# Sni445 recruits box C/D snoRNPs snR4 and snR45 to guide ribosomal RNA acetylation by Kre33

**DOI:** 10.1101/2025.08.22.671742

**Authors:** Jutta Hafner, Ingrid Zierler, Hussein Hamze, Sébastien Favre, Matthias Thoms, Natalia Kunowska, Sarah Rimser, Benjamin Albert, Tomas Caetano, Marion Aguirrebengoa, Roland Beckmann, Ulrich Stelzl, Dieter Kressler, Anthony K. Henras, Brigitte Pertschy

**Affiliations:** Institute of Molecular Biosciences, University of Graz, 8010 Graz, Austria; Molecular Cellular and Developmental Biology Unit (MCD), Centre de Biologie Intégrative (CBI), Université de Toulouse, CNRS, UPS, 31062, Toulouse, France; Unit of Biochemistry, Department of Biology, University of Fribourg, 1700 Fribourg, Switzerland; Gene Center, University of Munich, 81377 Munich, Germany; Institute of Pharmaceutical Sciences, Pharmaceutical Chemistry, University of Graz, 8010 Graz, Austria; BigA Core Facility, Centre de Biologie Intégrative (CBI), Université de Toulouse, 31062 Toulouse, France; BioTechMed-Graz, Graz, Austria; Biotech Research and Innovation Center, Faculty of Health and Medical Sciences, University of Copenhagen, 2200 Copenhagen, Denmark

**Keywords:** box C/D snoRNP, snR4, snR45, Kre33, cytidine acetylation, ribosome biogenesis, yeast

## Abstract

Eukaryotic ribosome synthesis is a highly complex, multistep process that is best characterized in the yeast *Saccharomyces cerevisiae*. It is orchestrated by over 200 ribosome assembly factors and 75 small nucleolar ribonucleoproteins (snoRNPs), which guide site-specific chemical modifications of precursor ribosomal RNA (pre-rRNA). While canonical box C/D snoRNPs direct 2’-O-methylation, the atypical box C/D snoRNPs snR4 and snR45 mediate acetylation of 18S rRNA residues C1280 and C1773, respectively, catalyzed by the acetyltransferase Kre33.

Here, we identify and characterize Ynl050c/Sni445 as a novel ribosome assembly factor and previously unrecognized auxiliary component of the snR4 and snR45 box C/D snoRNPs. Sni445 associates with snR4 and snR45 in their free form and is required for their stable incorporation into 90S pre-ribosomes. Genetic interactions link Sni445 and the snR4 and snR45 snoRNAs to ribosomal proteins Rps20 (uS10) and Rps14 (uS11), which are positioned near the respective acetylation sites in the 40S subunit. Moreover, Sni445 physically interacts with Kre33 within the 90S pre-ribosome, and its absence abolishes acetylation of C1280 and C1773.

Our findings suggest that Sni445 facilitates the recruitment of snR4 and snR45 snoRNPs to 90S particles and might promote their interaction with Kre33, thereby enabling the site-specific acetylation of 18S rRNA by Kre33.

## Introduction

Ribosomes are essential macromolecular machines that translate messenger RNA (mRNA) into proteins in all domains of life. In eukaryotes, ribosomes are composed of two subunits: the small 40S subunit and the large 60S subunit. The 40S ribosomal subunit contains the 18S ribosomal RNA (rRNA) and 33 ribosomal proteins (r-proteins). The 60S subunit consists of three rRNA species, 5S, 5.8S, and 25S rRNA in the model eukaryote *Saccharomyces cerevisiae* (yeast), and 5S, 5.8S, and 28S rRNA in humans, as well as 46 r-proteins in yeast and 47 in humans (1,2). The rRNA undergoes extensive modification, primarily through 2’-O-methylation and pseudouridylation. Other modifications, such as base methylation and acetylation are also introduced. Most of these modifications are introduced during the early steps of ribosome biogenesis (3,4).

Ribosome biogenesis is a conserved multi-step process that has been best characterized in yeast. More than 200 assembly factors transiently associate with pre-ribosomal particles to facilitate their maturation. This process begins in the nucleolus, where RNA polymerase I transcribes the 35S pre-rRNA, a precursor containing the sequences for the mature 18S, 5.8S and 25S rRNAs. These are flanked by external transcribed spacers (5’ ETS and 3’ ETS) and separated by internal transcribed spacers (ITS1 and ITS2). Spacer elements are then successively removed through endo- and exonucleolytic processing events. Already co-transcriptionally, the earliest r-proteins and assembly factors associate with the pre-rRNA to form the 90S pre-ribosome. Cleavage at site A_2_ within ITS1 separates precursors for the 40S and 60S subunits, which then undergo independent maturation pathways in the nucleolus and nucleoplasm, followed by nuclear export and final cytoplasmic maturation, culminating in the production of functional ribosomes (reviewed in (5–10)).

Most rRNA modifications are introduced by snoRNPs, each comprising a small nucleolar RNA (snoRNA) and a set of four core proteins. Through base-pairing, snoRNAs guide the snoRNPs to their target sites on the rRNA, where the catalytic subunit introduces the modifications. There are two classes of snoRNPs: box C/D snoRNPs that mediate 2’-O-methylation and box H/ACA snoRNPs that catalyze pseudouridylation. In yeast, box C/D snoRNPs include the core proteins Nop56, Nop58, Snu13 and the methyltransferase Nop1. Of the 46 box C/D snoRNAs expressed in yeast, 42 function in 2’-O-methylation, whereas the remaining four, i.e. U3, snR190, snR4 and snR45, do not guide methylation. U3 and snR190 play structural roles in ribosome maturation, while snR4 and snR45 guide the acetylation of cytidines C1280 and C1773 in the 18S rRNA (3,4,11–14). These modifications are introduced by the conserved Kre33 (NAT10 in humans), an acetyltransferase that also catalyzes tRNA modification with the help of its adaptor protein Tan1 and, moreover, fulfills a structural role within the 90S pre-ribosome (11,15–17).

Here, we introduce a novel ribosome assembly factor Sni445, functioning as an accessory protein component of the snR4 and snR45 acetylation guide snoRNPs. Sni445 is functionally connected to the r-proteins Rps20 (uS10^1^) and Rps14 (uS11), which bind in proximity to the snR4 and snR45 target sites. Sni445 binds to the free snoRNPs and is required for their docking to 90S pre-ribosomal particles. There, Sni445 interacts with the acetyltransferase Kre33, likely stabilizing the contact between Kre33 and snR4/snR45, thereby facilitating 18S rRNA acetylation.

## Material and Methods

### Yeast strain and plasmid construction

All *S. cerevisiae* strains that were used in this study are listed in Supplementary Table S1 and were generated by using established methods for chromosomal deletion and tagging (19,20). To introduce the *kre33.R637A* allele into the chromosome, a knock-in strategy based on homologous recombination was employed. For this purpose, a plasmid was constructed containing a recombination cassette composed of: (i) the 3’-terminal 1800 nucleotides of either the *KRE33* wild-type or *kre33.R637A* allele, (ii) 150 bp of downstream sequence corresponding to the *KRE33* terminator, (iii) a hygromycin resistance cassette, and (iv) chromosomal sequence spanning nucleotides 226 to 683 downstream of the *KRE33* open reading frame. The recombination cassette was PCR-amplified using the plasmid as a template, and the resulting linear PCR product was transformed into *RPS20* shuffle and *RPS14* shuffle strains. Hygromycin-resistant clones were selected on drug-containing plates, and correct genomic integration was confirmed by colony PCR and subsequent sequencing.

All plasmids generated in this study are listed in Supplementary Table S2 and were constructed using well-established molecular cloning methods using restriction enzymes. All DNA fragments that were generated by PCR were confirmed by sequencing.

### Microscopy

Yeast cells expressing C-terminally GFP-tagged Sni445 and the nucleolar marker protein Nop58-RedStar2 from the genomic locus were examined under the fluorescence microscope in logarithmic growth phase. For the Anchor Away technique (21), yeast cells genomically expressing the Sni445-FRB-GFP fusion protein were grown in logarithmic growth phase and were then incubated for up to 120 min with 1 µg/ml rapamycin in the absence or presence of 10 µg/ml cycloheximide. After various times of incubation, cells were visualized under the fluorescence microscope and compared to untreated cells. Both microscopy experiments were performed using a Leica DM6 B microscope with GFP or RHOD ET filters, a × 100/1.4 Plan APO objective and a DFC 9000 GT camera and the LasX software.

### Yeast two-hybrid (Y2H) assays

Yeast two-hybrid (Y2H) assays were performed to analyze protein-protein interactions. For this purpose, the Y2H reporter strain PJ69-4A (22) was transformed with two different plasmids, one plasmid encoding the bait protein fused N-terminally to the DNA-binding domain of the Gal4 transcription factor (abbreviated as BDN or G4BDN, with a c-myc tag and a *TRP1* marker used for selection) and the other plasmid encoding the prey protein fused C-terminally to the transcription activation domain of the Gal4 transcription factor (abbreviated as ADC or G4ADC, with a HA-tag and a *LEU2* marker used for selection). As controls, plasmids carrying just the DNA-binding domain of the Gal4 transcription factor (BDN-empty) or just the activation domain of the Gal4 transcription factor (ADC-empty) were used. After transformation, yeast cells were spotted in 10-fold serial dilutions starting with OD_600_ 0.75 on plates lacking tryptophan and leucine (SDC -Leu -Trp, serving as control plates); plates lacking tryptophan, leucine and histidine (SDC -Trp -Leu -His; growth on these plates indicates activation of the *HIS3* reporter gene, indicating a weak protein-protein interaction) or plates lacking tryptophan, leucine and adenine (SDC -Trp, -Leu, -Ade, growth on these plates indicates activation of the *ADE2* reporter gene, indicating a strong protein-protein interaction). Plates were incubated for 3 days at 30 °C.

### TurboID-based proximity labeling

The plasmid expressing C-terminally TurboID-tagged Sni445 under the control of the copper-inducible *CUP1* promoter was transformed into a wild-type yeast strain (YDK11-5A). The proximity-labelling experiment, mass spectrometry, data analysis and graphical representation were carried out as previously described (23). Raw and processed data are listed in Supplementary table S3.

### Genetic interaction tests

To analyze genetic interactions, *SNI445* was knocked out in *rps24a*Δ or *rps24b*Δ cells, or in *RPS2*, *RPS14,* or *RPS20* shuffle strains, carrying *URA3* plasmids containing the respective wild-type genes. Moreover, *SNR4* or *SNR45* were knocked out or the *kre33.R637A* allele was chromosomally integrated in *RPS14* or *RPS20* shuffle strains. For examination of genetic interactions between *SNI445* and *RPS24*, double and single mutants, as well as the wild-type strain were spotted in serial dilutions starting with an OD_600_ of 0.75 on synthetic dextrose complete (SDC) plates. For the experiments with shuffle strains, cells were transformed with *LEU2* plasmids carrying either wild-type or different alleles of *RPS20* and *RPS14* or with *TRP1* plasmids carrying *RPS2* or the *rps2-1* allele. After transformation, cells were streaked on SDC plates (SDC + all) containing 1 g/L 5-FOA (Thermo Scientific) to select for cells that lost the *URA3* plasmid carrying the respective wild-type gene. Subsequently, cells were spotted on SDC-Leu (*RPS20*, *RPS14*) or SDC-Trp (*RPS2*) agar plates in 10-fold serial dilutions starting with OD_600_ 0.75. Plates were incubated for 2-3 days at 25°C, 30°C and 37°C. Cells which were not able to grow on 5-FOA containing agar (*rps14a.R136A sni445*Δ, *rps14a.R136A* Δs*nr45*, *rps14a.R136A kre33.R637A*) and respective controls were spotted on SDC -Leu plates and SDC plates containing 5-FOA in serial dilutions starting with OD_600_ 2 and incubated at 30 °C for 3-6 days.

### CRAC experiment and analysis

For CRAC experiments, a yeast strain expressing from the chromosomal locus a Sni445 fusion protein bearing a C-terminal HTP-tag (His6 tag-TEV protease cleavage site-Protein A tag) and an untagged wild-type strain (negative control) were used. The CRAC experiment was performed as previously described in (24) and (25). The cDNA samples were sent for Illumina NextSeq2000 deep sequencing (EpiRNA-Seq facility, CNRS, Université de Lorraine, INSERM).

CRAC-seq data processing was performed using the pyCRAC toolkit (version 1.5.1) (https://sandergranneman.bio.ed.ac.uk/pycrac-software), a suite of Python-based tools specifically designed for the analysis of Crosslinking and Analysis of cDNAs (CRAC) data. The analysis pipeline was implemented using Snakemake (version 8.5.5), ensuring workflow reproducibility, modularity, and scalability. Each step of the pipeline was encapsulated within dedicated conda environments, guaranteeing consistent software dependencies across executions.

- Raw reads were first demultiplexed using pyCRAC’s ‘pyBarcodeFilter.py’ with 1 mismatch allowed.
- Raw reads were quality-checked using FastQC (v0.12.1) and aligned to the _Saccharomyces cerevisiae_ pyCRAC’s reference genome (Saccharomyces_cerevisiae.EF2.59.1.0.fa) using Novoalign (v3.09.00) with the ‘-r Random’ parameter to randomly assign multimapping reads. Both ‘.sam’ and ‘.novò output files were retained to support downstream analyses: ‘.sam’ files were used to generate CPM-normalized BigWig coverage tracks using **samtools** (v1.14) for sorting and indexing, and **deepTools** (v3.4.3) for coverage computation (’bamCoveragè); ‘.novò files were used as input for pyCRAC’s internal tools requiring alignment metadata.
- The pyCRAC’s reference GTF annotation file (Saccharomyces_cerevisiae.EF2.59.1.3.gtf) was first checked using PyCRAC’s ‘pyCheckGTFfile.py’. Gene name information was extracted using PyCRAC’s ‘pyGetGeneNamesFromGTF.py’, and a tabular genome file required by pyCRAC was generated from the reference FASTA using a custom Python script.
- PyCRAC’s pyReadCounters.py was used to quantify the number of crosslinking events per transcript. The tool was run without the ‘--blocks’ option, using various values for the ‘--readLength’ parameter, in this study we generated data at 30 and 1000 bp read lengths. All hits were analyzed without limitation applied.
- Pileup profiles of crosslink events were generated using PyCRAC’s pypileup.py. This tool was run with default parameters with pyreadCounter output gtf as input.

All configuration files, custom scripts, and conda environment definitions used to run the workflow are available upon request, enabling full reproducibility of the analysis.

NGS analysis files of raw and processed data were deposited in the Gene Expression Omnibus database under the accession number GSE299803 (For review only: secure token: ebkbqmkyjjclfon).

### Tandem affinity purification (TAP)

Cells expressing chromosomally C-terminally TAP-tagged (calmodulin binding peptide-TEV protease cleavage site-protein A tag) fusion proteins were used for TAP purifications. To this end, cells were grown in 2 L YPD to an OD_600_ of 1.8, harvested and frozen at -20 °C. Pellets were suspended in lysis buffer (50 mM Tris-HCl pH 7.5, 100 mM NaCl, 1.5 mM MgCl_2_, 0.075 % NP-40, 1 mM dithiothreitol (DTT, Roth) and 1x Protease Inhibitor Mix FY (Serva)) and lysed by mechanical disruption using glass beads. After a centrifugation step, the cleared cell lysates were incubated with 300 µl IgG SepharoseTM 6 Fast Flow (GE Healthcare) for 1.5 h at 4°C. IgG beads were then transferred into Mobicol-columns (MoBiTec) and washed with 10 ml lysis buffer. After washing beads, 300 µl of buffer containing TEV protease was added and bound proteins were eluted for 70 min at room temperature. CaCl_2_ was added to TEV eluates to a final concentration of 2 mM and the TEV eluate was then incubated with 65 µl Calmodulin Sepharose^TM^ 4B (GE Healthcare) for 1 h at 4°C. After washing beads with 5 ml buffer containing 2 mM CaCl_2_, elution of bound proteins was performed using 40 µl of elution buffer containing 5 mM EGTA for 20 min at room temperature. Eluates were then used for RNA isolation or samples were subjected to mass spectrometry (MS) analysis. Final samples were further analyzed on NuPAGE^TM^ 4-12% Bis-Tris gels followed by NOVEX^®^ Colloidal Blue Staining Kit (Thermo Fisher) staining.

### FLAG purification

Cells expressing C-terminally FLAG-tagged Sni445 from the genomic locus were grown in 2 L YPD to an OD_600_ of 1.8. Cells were harvested and frozen at – 20 °C. Pellets were suspended in lysis buffer (50 mM Tris-HCl pH 7.5, 100 mM NaCl, 1.5 mM MgCl_2_, 0.075 % NP-40, 1 mM DTT and 1x Protease Inhibitor Mix FY). Cells were then lysed by mechanical disruption using glass beads. After a centrifugation step, the cleared lysates were incubated with 300 µl Anti-FLAG^®^ M2 Affinity Gel (Sigma Aldrich) for 1 h at 4 °C. Beads were then transferred to Mobicol-columns (MoBiTec) and washed with 10 ml buffer. Elution of bound proteins was performed using buffer containing 100 µg/ml FLAG-peptide (Sigma Aldrich) for 1 h at 4°C. Eluates were then used for RNA isolation or samples were analyzed by MS analysis. Final samples were further analyzed on NuPAGE^TM^ 4-12% Bis-Tris gels followed by NOVEX^®^ Colloidal Blue Staining Kit (Thermo Fisher) staining.

### Split tag purification of Sni445-TAP Nop58-FLAG

Cells were grown to an OD_600_ of ∼ 3, harvested by centrifugation and flash frozen in liquid nitrogen and stored at -80°C. Cells were lysed using a SPEX 6970EFM Freezer/Mill. Cell powder was resuspended in buffer containing 60 mM Tris-HCl pH 7.5, 50 mM NaCl, 40 mM KCl, 5 mM MgCl_2_, 1mM DTT supplemented with 5% glycerol, 0.1% NP-40 and 1x EDTA-free protease inhibitor (Roche). The lysate was cleared by two successive centrifugation steps: first at 4,000 rpm for 15 minutes (Eppendorf 5810 R), and second at 17,500 rpm for 25 minutes (Sorvall LYNX 6000). Both steps were carried out at 4°C. The supernatant was incubated with pre-equilibrated IgG Sepharose^TM^ 6 Fast Flow affinity resin and incubated at 4 °C on a turning wheel for 90 min. IgG beads were collected by centrifugation and after a batch wash transferred to a Mobicol column (MoBiTec) and washed with additional 10 ml lysis buffer by gravity flow. Bound proteins were eluted from the beads by the addition of homemade TEV protease by incubating for 90 min at 16°C. Anti-FLAG M2 agarose beads (Sigma-Aldrich) were added to the eluate and incubated for 60 min at 4°C. The beads were washed with buffer supplemented with 0.01% NP-40 and 2% glycerol and transferred to a Mobicol column (MoBiTec) for a final wash with 10 ml buffer containing 0.05% Nikkol and 2% glycerol. Samples were eluted by the addition of 3x Flag peptide (Sigma-Aldrich, final concentration 250 µg/ml) for 1 h at 4°C and analysed on a 4-12% polyacrylamide gel (NuPAGE, Invitrogen) stained with colloidal Coomassie. Copurifying proteins were analysed by mass spectrometry.

### Mass Spectrometry proteomics

#### Sample processing

Eluates of the Sni445-FLAG and Sni445-TAP purification were processed using S-Trap™ micro columns (ProtiFi, Cat# C02-micro-80) following the high recovery protocol recommended by the manufacturer and using trypsin (Pierce, Cat# 90059) as the carrier protein. After elution from the columns, the samples were lyophilized and resuspended in 12 µl 0.1% formic acid; then 1 µl thereof was used per MS injection.

#### Mass spectrometry

Tryptic peptide samples were analyzed on an UltiMate 3000 RSLC (Thermo Scientific) coupled to a TimsTOF PRO (Bruker) mass spectrometer. Peptides were separated on a reversed-phase C18 Aurora column (25 cm × 75 µm) with an integrated Captive Spray Emitter (IonOpticks) at a column temperature of 50°C and a flow rate of 300 nl/min. Mobile phases were A, 0.1% (v/v) formic acid in water, and B, 0.1% (v/v) formic acid in acetonitrile, respectively. Fraction B was linearly increased from 2% to 25% in a 90-min gradient, then increased to 40% for 10 min, and finally further increased to 80% for 10 min, followed by re-equilibration. The column was washed between runs with 50% buffer B for 3 min, 80% buffer B for 3 min, and re-equilibration with 2% buffer B for 5 min. The spectra were recorded in data-independent acquisition (DIA) mode.

#### Data processing

The DIA data were quantified with DIA-NN v.1.9 using a synthetic fasta library computed from the UniProt *S. cerevisiae* protein database file (reviewed; 2022-03-20), with default settings. MS2 and MS1 mass accuracies were set to 20 ppm and scan window size was set to 9. Output was filtered at 0.01 FDR.

#### Data analysis

The Label-Free Quantification (LFQ) output table generated in DIA-NN was filtered, and only the proteins detected in at least 3 out of 6 MS runs for any of the pulldowns were kept. The LFQ values were then log_2_-transformed, z-score-and bait-normalized, and the values for each biological replicate were averaged across the two MS runs. Next, the mean value for the three biological replicates was calculated and the missing values were imputed using Perseus style imputation with random values drawn from a normal distribution downshifted by 1.8 standard deviation with a width of 0.3 for each sample. Finally, the difference between averaged log_2_ intensity values of Sni445-TAP and Sni445-FLAG and the average normalized label-free quantification from all experiments were calculated.

The MS proteomics raw data together with the processing files were deposited to the ProteomeXchange Consortium using the PRIDE partner repository (https://www.ebi.ac.uk/pride/) with the dataset identifier PXD065447. Reviewers can access the dataset by log in to the PRIDE website using the following details: Project accession: PXD065447; Token: vzPAtkd6lx9E.

### Northern Blotting

For RNA isolation, crude extract and final eluate from TAP or FLAG purifications were adjusted to a final volume of 600 µl with the respective buffers. RNA was extracted in two rounds of phenol-chloroform-isoamyl alcohol extraction (25:24:1) followed by one round of chloroform-isoamyl alcohol (24:1) extraction. RNA was then precipitated by adding 1/10 volume of 3 M sodium acetate pH 5.2, 2.5 volumes of 100 % ethanol and 1 µl of GylcoBlue^TM^ (Invitrogen). After precipitation over night at -20 °C and pelleting of precipitated RNA by centrifugation, RNA pellets were dissolved in nuclease-free water. RNA samples were then loaded on NuPAGE^TM^ 6% TBE-Urea gels (Thermo Fisher) followed by blotting onto Hybond N^+^ nylon membranes (Amersham Biosciences) using 0.5X TBE buffer at 300 mA. To detect different snoRNA species, oligonucleotides complementary to the snoRNAs of interest were 5’-^32^P-labeled using T4 polynucleotide kinase and γ-[^32^P]ATP (185TBq (5000Ci)/mmol (Hartmann analytics). To enhance signals, several oligonucleotide probes complementary to different regions of the same snoRNAs were combined. The following oligonucleotides were used: snR4-1: 5’- CATCGACCCAGGAAAGCATCTTACAC -3’, snR4-2: 5’- CTAGAGTTATTTTAAAACAC -3’, snR4-3: 5’- CTATAACCTATCCTCATCGACTG -3’, snR45-1 : 5’- CAGATCGCTCCGAGAAGAATTG -3’, snR45-2: 5’- GCGCAGGAACCGCTATCTCC -3’, snR45- 3: 5’- CATTCTTAAGAATGTAACAAGATCAATGGG -3’, U3: 5’- GGATTGCGGACCAAGCTAA -3’, snR10-1: 5’- CCTTGTCGTCATGGTCGAATCG -3’, snR10-2: 5’- TCCTTGCAACGGTCCTCATC -3’, snR35-1: 5’- GAAGCCTAAACTTCCCTCAATTTCCTACAC - 3’, snR35-2: 5’- CCAAAAGAGACTCGATATAAACAACACGG -3’, snR35-3: 5’- GCATGTCTGTCCTACCAGCCCTTGCATAGGCG -3’. Hybridization with the radiolabeled oligonucleotides was performed at 37°C overnight in buffer containing 0.5 M Na_2_HPO_4_, pH 7.2, 7% SDS, and 1 mM EDTA. After three subsequent washing steps with a buffer containing 40 mM Na2HPO4, pH 7.2, 1% SDS, signals were detected by exposing X-ray films. Membranes were regenerated by washing in 1% SDS prior to hybridization.

### Reverse transcription and misincorporation analysis of ac4C in 18S RNA

For detection of acetylated cytidines, we adapted the assay developed in (26). Cells were grown in 20 mL YPD at 30 °C to an OD_600_ of 0.7-0.9 and 4 OD_600_ units were harvested. Cells were resuspended in 200 µL lysis buffer containing 40 mM Tris-HCl pH 7.5, 40 mM EDTA, and 2% SDS and mechanically lysed by vigorous shaking with 200 µL glass beads (0.5 mm diameter) for three minutes. RNA was extracted from the lysate by three extraction steps with phenol-chloroform-isoamyl alcohol (25:24:1) and one extraction step with chloroform-isoamyl alcohol (24:1). RNA was then precipitated by addition of 1/10 volume of 3 M sodium acetate (pH 5.2), 2.5 volumes of 100% ethanol and 1 µL GlycoBlue^TM^ (Invitrogen), and after drying, RNA was dissolved in 30 µL nuclease-free water.

For chemical reduction of ac4C in 18S rRNA, 1 µg of the isolated RNA was either incubated with H_2_O or 100 mM sodium borohydride (NaBH_4_) in a final reaction volume of 100 µL. Samples were incubated at 37°C for one hour, quenched with 15 µL HCl (1 M) and neutralized by addition of 15 µL Tris-HCl buffer (1 M, pH 8.0) in a final volume of 200 µL. RNA precipitation was performed by addition of 1/10 volume of 3 M sodium acetate (pH 5.2), 2.5 volumes of 100% ethanol, followed by centrifugation at 4°C, 15,000 rpm for 15 min. The RNA pellet was washed with 70% EtOH, and after drying, dissolved in 10 µL nuclease-free water.

For reverse transcription, 200 pg RNA from individual treatment reactions were incubated with 4 pmols of the respective primers (18S_h34_rev (5’-GGTTAAGGTCTCGTTCGTTATCG-3’) for analysis of acetylation in helix h34 or 18S_h45_rev (5’-TAATGATCCTTCCGCAGGTTCACCTAC-3’) for analysis of acetylation in helix h45), 10 nmol dNTP mix (NEB), and nuclease-free water in a final reaction volume of 10 µL, incubated at 65°C for 5 minutes and promptly transferred on ice. 1x Induro RT Reaction Buffer (NEB), 8 units Murine RNase Inhibitor (NEB) and 200 units Induro Reverse Transcriptase (NEB) were added in a final volume of 20 µL. Samples were incubated at 55°C for 10 min, followed by inactivation at 95°C for 1 minute.

2 µL of the cDNA products were used as templates in a 50 µL PCR reaction with 1x Q5 reaction buffer (NEB), 25 pmols of forward primers (18S_h34_fwd: 5’- AAGGAATTGACGGAAGGGC-3’ or 18S_h45_fwd: 5’- CGTCGCTAGTACCGATTGAATGGCTTAG -3’) and 25 pmols of the respective reverse primers used for the reverse transcription (18S_h34_rev or 18S_h45_rev, respectively), 10 nmol dNTP mix, and 2.5 units Q5 Polymerase (NEB). PCR was performed with an initial denaturation at 98 °C for 30 sec; followed by 34 cycles of denaturation at 98 °C for 10 sec, annealing at 66 °C (helix 34) or 71°C (helix 45) for 30 sec, elongation at 72 °C for 30 sec; followed by final elongation at 72°C for 2 min.

For Sanger sequencing, 5 µL of each PCR product was cleaned up using the Exo-CIP Rapid PCR Cleanup Kit (NEB) and sequenced using the forward PCR primer. Processed sequencing traces were viewed using Chromas.

## Results

### Ynl050c/Sni445 physically and genetically interacts with small subunit r-proteins

To identify novel interaction partners of small ribosomal subunit r-proteins, we previously conducted a tandem affinity purification (TAP)-based screen (27). Using TAP-tagged versions of Rps10 (eS10), Rps12 (eS12), Rps13 (uS15), Rps14 (uS11), Rps18 (uS13), Rps19 (eS19), Rps20 (uS10), Rps24 (eS24), Rps31 (eS31), which bind different domains of the 40S subunit (Figure 1A) - as baits, we identified the uncharacterized protein Ynl050c, which we subsequently named Sni445 (see below), as a potential interactor (27). Sni445 (Ynl050c) is predicted to be an unstructured protein with a negatively charged N- and a positively charged C-terminal part (Figure 1B). The middle part of Sni445 (amino acids 142-241) harbors an a-helical patch with no defined domain structure while the rest of the protein is highly disordered.

**Figure 1:**
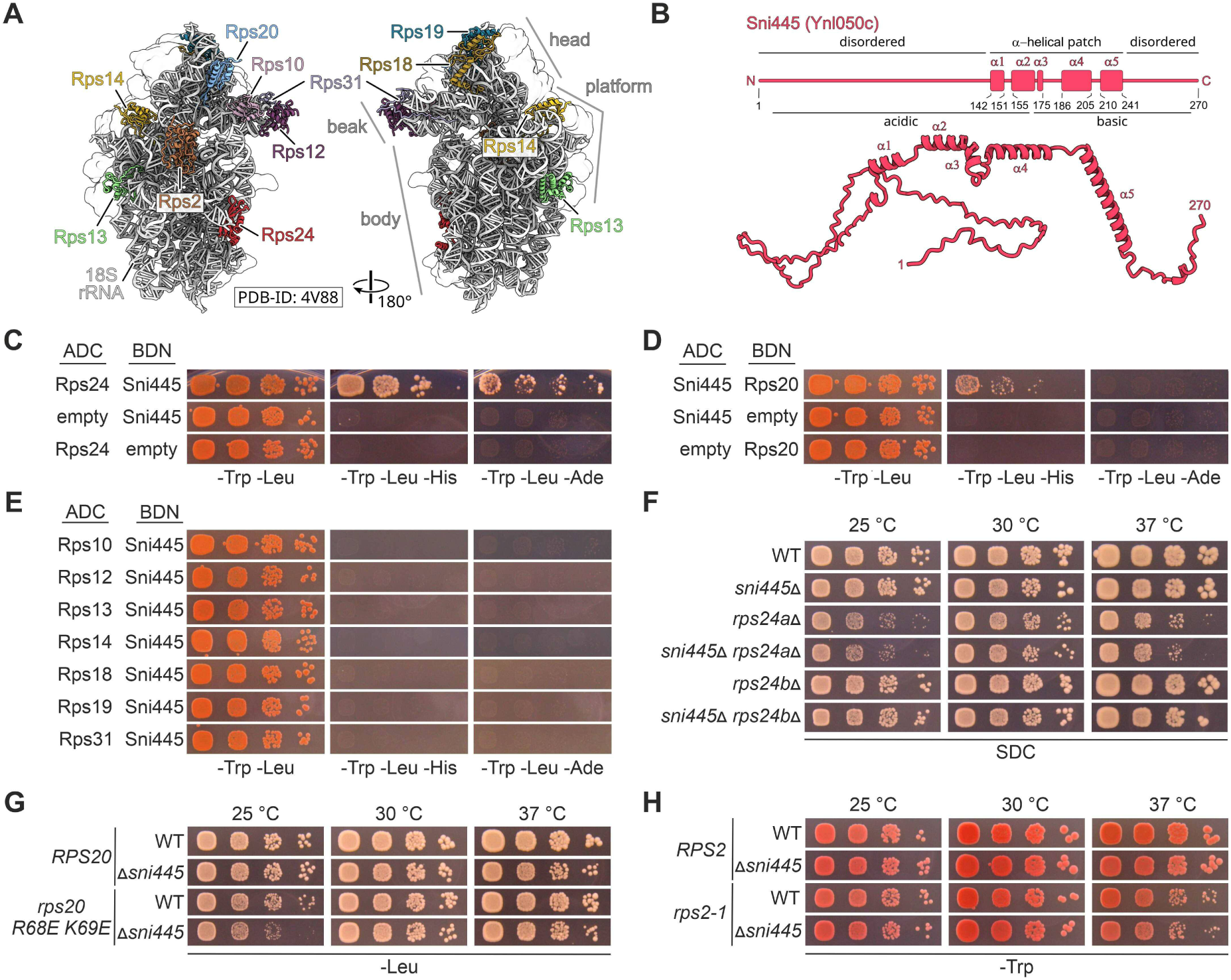
Ynl050c/Sni445 physically and genetically interacts with r-proteins. (A) Structure of the 40S subunit (PDB-ID: 4V88; (1)). The 18S rRNA is shown in gray and r-proteins analyzed in the study are colored and labeled. (B) Secondary structure (upper panel) and AlphaFold (52) prediction (lower panel) of Sni445 (Ynl050c). (C–E) Sni445 physically interacts with Rps24 and Rps20. Y2H interaction assays between Sni445 and r-proteins Rps24 (C), Rps20 (D), and Rps10, Rps12, Rps13, Rps14, Rps18, Rps19, and Rps31 (E). Proteins were fused either C-terminally to the Gal4 activation domain (ADC) or N-terminally to the Gal4 DNA-binding domain (BDN). Combinations with empty vectors were tested as negative controls. Cells were spotted in serial dilutions on SDC-Trp-Leu plates (growth control), SDC-Trp-Leu-His plates (growth indicates weak interaction), and SDC-Trp-Leu-Ade plates (growth indicates strong interaction) and incubated for three days at 30°C. (F) No enhanced growth defect in *rps24aΔ sni445Δ* or *rps24bΔ sni445Δ* double mutants compared to the respective single mutants. Wild-type and mutant cells were spotted on SDC plates and incubated at 25°C, 30°C, or 37°C for three days. (G) Genetic enhancement of the growth defect of the *rps20*.*R68E/K69E* mutant upon deletion of *SNI445*. Yeast cells (*rps20*Δ) carrying *LEU2* plasmids with either wild-type *RPS20* or the mutant *rps20.R68E/K69E* allele, either in a *SNI445* wild-type background (WT) or combined with a *SNI445* knockout (*sni445*Δ), were spotted in serial dilutions on SDC-Leu plates and incubated at 25°C for three days, or 30°C or 37°C for two days. (H) No genetic interaction between *SNI445* and *RPS2*. Yeast cells (*rps2*Δ) carrying *TRP1* plasmids with either wild-type *RPS2* or the mutant *rps2-1* allele, either in a *SNI445* wild-type background (WT) or combined with a *SNI445* knockout (*sni445*Δ), were spotted in serial dilutions on SDC-Trp plates and incubated at 25°C, 30°C, or 37°C for three days.

To test whether Sni445 physically interacts with any of these r-proteins, we performed yeast two-hybrid (Y2H) assays. These revealed a relatively strong interaction between Sni445 and Rps24 (Figure 1C), and a weaker interaction with Rps20 (Figure 1D). No Y2H interactions were detected with Rps10, Rps12, Rps13, Rps14, Rps18, Rps19, or Rps31 (Figure 1E). Based on the rationale that functional connections between genes are often revealed by genetic interactions, such as enhanced growth defects when mutations in both genes are combined, we next investigated whether *SNI445* genetically interacts with either *RPS24* or *RPS20*. Since no *rps24* point mutants were available, we took advantage of the fact that Rps24 is encoded by two paralogous genes in yeast, *RPS24A* and *RPS24B*. We generated *sni445*Δ *rps24a*Δ and *sni445*Δ *rps24b*Δ double deletion mutants and compared their growth with the one of the respective single deletion mutants. However, combining the deletion of *SNI445*, which on its own did not affect growth, with either of the two *rps24* null alleles did not result in a synergistic growth defect (Figure 1F), suggesting the absence of genetic interactions in this context. In contrast, a previously generated *rps20.R68E/K69E* allele, which substitutes two basic residues, arginine and lysine, in an unstructured loop with glutamates, resulting in defects in early cytoplasmic pre-40S maturation (28), showed a mild growth defect that was exacerbated by deletion of *SNI445*, particularly at 25°C (Figure 1G). This indicates a genetic interaction between *SNI445* and *RPS20*. In contrast, no synthetic growth defects were observed in a control experiment combining the unrelated *rps2-1* allele (27) with *sni445*Δ (Figure 1H).

In summary, our data indicate that Sni445 may physically interact with both Rps24 and Rps20 and that Sni445 and Rps20 are functionally connected by a genetic interaction.

### Sni445 is a ribosome assembly factor

The interactions of Sni445 with r-proteins could either occur during translation or ribosome assembly. To distinguish between these possibilities, we analyzed the subcellular localization of Sni445, based on the rationale that translation occurs in the cytoplasm, whereas ribosome biogenesis predominantly takes place in the nucleus. Fluorescence microcopy revealed that Sni445-GFP co-localizes with the nucleolar marker Nop58-RedStar2, indicating that its steady-state localization is in the nucleolus (Figure 2A), suggesting a role for Sni445 in early steps of ribosome assembly.

**Figure 2:**
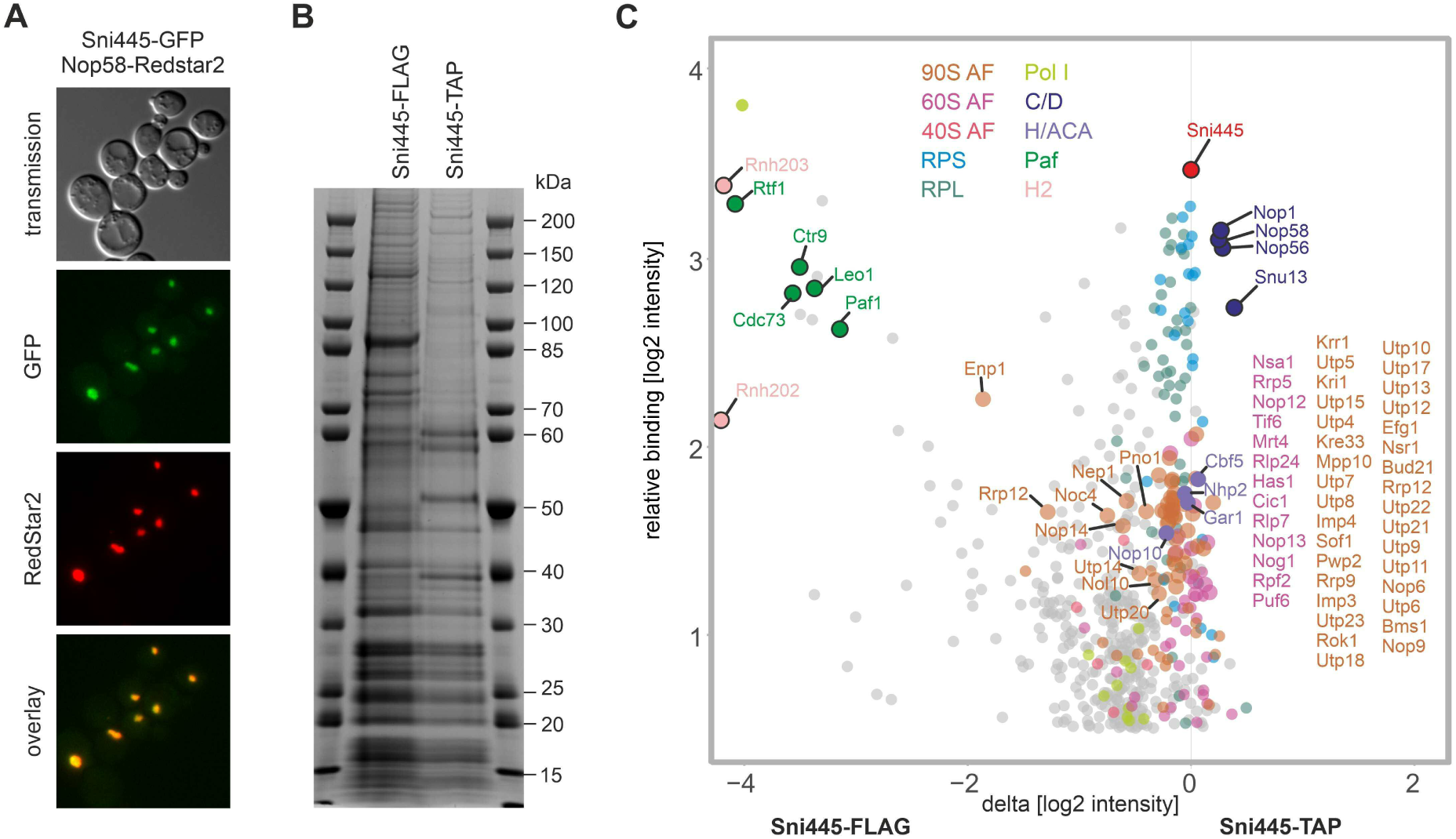
Sni445 is a novel ribosome assembly factor. (A) Fluorescence microscopy reveals that Sni445-GFP localizes to the nucleolus. Nop58-RedStar2 was used as a nucleolar marker. Overlay images show co-localization of Nop58-RedStar2 with Sni445-GFP. (B) Sni445-TAP and Sni445-FLAG eluates were analyzed by SDS-PAGE and Coomassie blue staining. (C) Sni445 co-purifies ribosome assembly factors and box C/D snoRNA core proteins. Sni445-FLAG and Sni445-TAP eluates were analyzed by label free mass spectrometry in three biological replicates. z-score-normalized label-free quantification (LFQ) values were log2-transformed and visualized. X-axis (delta [log2 intensity]), difference of averaged log2 intensity values of ([Sni445-TAP]-[Sni445-FLAG]) normalized to the Sni445 protein. Y-axis (relative binding [log2 intensity]): averaged median-normalized label-free quantification from all experiments. Only specifically bound proteins with values >0.5 are shown (552 out of 2869). Proteins are annotated through color labels: Sni445 (red); box C/D proteins (C/D; 4, blue); box H/ACA proteins (H/ACA; 4, purple); Paf complex (Paf; 5, green); RNaseH2 (H2, light pink); polymerase I subunits (Pol I; 11 light yellow); 90S assembly factors (90S AF; 65, brick); 60S assembly factors (60S AF; 40, magenta); 40S assembly factors (40S AF; 6, pink); small subunit r-proteins (RPS; 23, cyan); large subunit r-proteins (RPL; 45, turquoise).

Several ribosome assembly factors show a nucleolar steady-state localization, but shuttle between the nucleus and cytoplasm (29,30). To assess whether Sni445-GFP is a shuttling protein or an exclusively nuclear protein, we employed the anchor-away technique (21). Sni445 was fused to FRB-GFP (FKBP12 -rapamycin-binding domain of human mTOR), in a rapamycin resistant *tor1-1* strain, in which the plasma membrane-located protein Pma1 was fused to FKBP12 (human FK506 binding protein). Upon rapamycin treatment, a ternary complex between FRB, FKBP12, and rapamycin is formed, leading to the sequestration of FRB-GFP tagged proteins at the plasma membrane if the protein enters the cytoplasm in the course of its functional cycle (21,29). Many known shuttling proteins relocalize to the plasma membrane within ∼15 minutes of rapamycin treatment (30). In contrast, Sni445-FRB-GFP remained nucleolar even after 30 minutes of rapamycin treatment (Supplementary Figure S1). A faint plasma membrane signal only became detectable after 60 minutes, likely due to de novo protein synthesis of Sni445-FRB-GFP. Supporting this, co-treatment with cycloheximide to block protein synthesis prevented plasma membrane localization, confirming that Sni445 does not shuttle and is an exclusively nucleolar protein (Supplementary Figure S1).

To determine the stage of ribosome assembly in which Sni445 functions, we analyzed its protein interactome. Sni445 was fused to either a C-terminal TAP- or FLAG-tag and affinity purified from exponentially grown yeast cells. SDS-PAGE followed by Coomassie blue staining revealed fewer protein bands in the TAP eluate than in the FLAG eluate (Figure 2B). This could reflect reduced contamination due to the two-step TAP purification or, alternatively, a disruption of some of the protein-protein interactions by the bulky TAP tag. Mass spectrometry showed that several proteins were preferentially co-enriched with Sni445-FLAG, suggesting that the large TAP tag might indeed disturb certain interactions. In particular, Rtf1, Ctr9, Leo1, and Paf1, all components of the polymerase-associated factor (Paf) complex, which is involved in RNA polymerase II transcription, 3’ end formation of mRNAs and snoRNAs (31–33), as well as RNA polymerase I transcription (34,35), co-purified with Sni445-FLAG (Figure 2C). Additionally, RNase H2 subunits Rnh202 (H2B subunit) and Rnh203 (H2C subunit) were also strongly co-purified (Figure 2C).

Importantly, multiple proteins co-purified with both Sni445-TAP and Sni445-FLAG, including r-proteins, 90S particle assembly factors (e.g., Krr1, Kri1, Utp5, Utp22), and pre-60S particle factors, such as Nsa1 and Nop12. Most notably, and consistent with a recent high-throughput study (28), we found all four core proteins of box C/D snoRNPs, Nop1, Nop56, Nop58, and Snu13, strongly enriched in both the Sni445-TAP and Sni445-FLAG eluates, while box H/ACA snoRNP core proteins (Cbf5, Nhp2, Gar1, and Nop10) were also detected, but were not particularly enriched (Figure 2C). Taken together, these findings indicate that Sni445 associates with early pre-ribosomal particles, and possibly engages with the Paf complex and with box C/D snoRNPs. We therefore conclude that Sni445 is involved in ribosome biogenesis, most likely in a function in the context of box C/D snoRNPs.

### Sni445 is a component of the snR4 and snR45 acetylation guide snoRNPs

Based on its co-purification with pre-ribosomal particles and box C/D snoRNP proteins, we speculated that Sni445 might interact with pre-ribosomal RNA and/or snoRNAs. To identify RNA sequences bound by Sni445, we performed RNA crosslinking and analysis of cDNA (CRAC) (36). Strikingly, ∼80 % of the sequence reads mapped to snoRNAs, with the vast majority corresponding to only two snoRNAs, the box C/D snoRNAs snR4 (∼47 % of all reads) and snR45 (∼32 % of all reads) (Figure 3A, Supplementary Figure S2). These two snoRNAs do not guide methylation, but instead direct acetylation of cytidines C1280 and C1773 in the 18S rRNA via the acetyltransferase Kre33. These findings prompted us to name the protein Sni445 (**<u>sn</u>**oRNA **i**nteractor of snR**<u>4</u>** and snR**<u>45</u>**).

**Figure 3:**
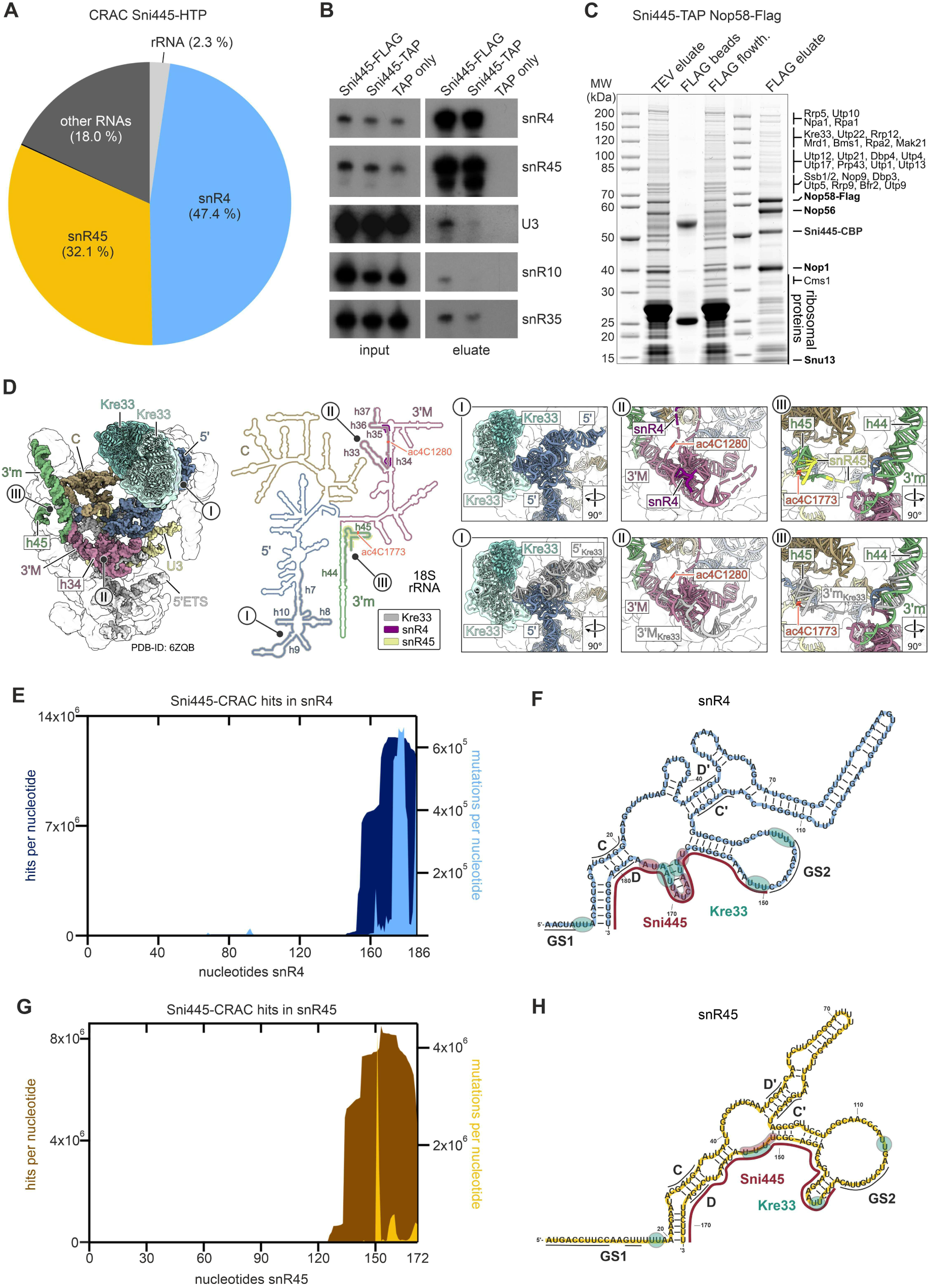
Sni445 associates with the acetylation guide snoRNAs snR4 and snR45. (A) CRAC analysis reveals that Sni445 associates with the box C/D snoRNAs snR4 (blue) and snR45 (dark yellow). Pie chart shows RNA species crosslinked with Sni445-HTP. See also Supplementary Figure S2 for comparison to the negative control.

(B) snR4 and snR45 are strongly co-enriched with Sni445. RNA was extracted from Sni445-TAP, Sni445-FLAG, or TAP only eluates. Northern blot analysis was performed on inputs and eluates using the indicated probes. The inputs and eluates were analyzed on the same gel and the same exposures are shown, allowing direct comparisons.

(C) Split-tag purification of complexes containing both Sni445 and Nop58. Complexes were first purified via Sni445-TAP followed by Nop58-FLAG, and analyzed by SDS-PAGE and Coomassie blue staining. The TEV eluate (all proteins co-purified with Sni445-TAP), the FLAG flowthrough (unbound fraction) and FLAG bead fraction (proteins not eluted by FLAG peptide) were loaded as controls. Proteins identified by mass spectrometry are indicated. (D) Structure of a 90S pre-ribosome in state B2 (PDB 6ZQB, (37)), shown alongside the secondary structure of 18S rRNA. The Kre33 dimer is highlighted in turquoise, and the structural domains of the 18S rRNA are color coded: 5’ domain in blue, central domain in brown, 3’ major domain in magenta, 3’ minor domain in green. The U3 snoRNA is shown in bright yellow. Kre33 crosslinking sites identified by CRAC (11) are indicated in the 3D and 2D structure (I for the crosslinking site in the 5’ domain, II for the site in helix h34, and III for the site in helix h45). Detailed views of these regions are provided in the zoom-in panels. See also Supplementary Figure S3 for a more detailed representation of the 18S rRNA secondary structure. (E) Distribution of sequence reads (dark blue) and mutations/deletions (light blue) in snR4 recovered in the Sni445-HTP CRAC experiment. The left y-axis shows read counts; the right y-axis shows mutation frequencies. Mutations arise from crosslink-induced reverse transcription errors and thus mark crosslinking sites. (F) Secondary structure of snR4 (blue) (11). snR4 contains two guide sequences (GS1 and GS2, black lines) that base-pair with 18S rRNA, and conserved box C/D and box C’/D’ motifs (black lines). Major sequencing hits from Sni445 CRAC are indicated in red lines, with the crosslinking sites indicated by mutations/deletions marked by red bubbles; major Kre33 crosslinking hits (11), are indicated by green bubbles. (G) Distribution of sequence reads (brown; left y-axis) and mutations (dark yellow; right y-axis) in snR45 from the Sni445-HTP CRAC experiment. (H) Predicted secondary structure of snR45 (yellow). snR45 harbors two guide sequences (GS1 and GS2, black lines) that base-pair with 18S rRNA and conserved box C/D and box C’/D’ motifs (black lines). Major sequencing hits from Sni445 CRAC are indicated in red lines, with crosslinking sites indicated by mutations/deletions marked by red bubbles; major Kre33 crosslinking hits (11), are indicated by turquoise bubbles.

To confirm this result using other means, we performed northern blotting analyses on eluates from Sni445-TAP and Sni445-FLAG purifications. We observed a strong enrichment of snR4 and snR45 compared to other snoRNAs, further confirming their association with Sni445 (Figure 3B, compare signals in inputs and eluates).

To further characterize the complex(es) containing Sni445 and the snR4/snR45 snoRNPs, we employed a split two-step affinity purification strategy. First, Sni445-TAP was purified, followed by a second purification step using the box C/D core protein Nop58 tagged with a FLAG tag as bait. This strategy yielded stochiometric amounts of Sni445 along with the canonical box C/D snoRNP core proteins Nop1, Nop56, and Nop58 (Figure 3C), indicating that Sni445 forms a stable complex with box C/D snoRNPs not bound to pre-ribosomal particles. In addition, several substoichiometric bands corresponding to 90S assembly factors, including Rrp5, Utp10, Utp22, Rrp12 and Kre33, as well as pre-60S assembly factors such as Npa1 and Mak21, were detected (Figure 3C), suggesting that a subset of the Sni445-containing box C/D complexes are associated with early pre-ribosomal particles.

Taken together, these data demonstrate that Sni445 is a bona fide component of the snR4 and snR45 snoRNPs, associating with them prior to their engagement with pre-ribosomal particles. The presence of 90S assembly factors in the purifications further suggests that Sni445-bound snoRNPs are also part of 90S pre-ribosomal particles.

snR4 and snR45 direct site-specific acetylation by base-pairing with distinct regions of the 18S rRNA. Specifically, snR4 targets helix h34 in the 3’ major domain, to guide acetylation at C1280 (II in Figure 3D (11)). snR45 pairs with helix h45 in the 3’ minor domain, near the 3’ end of the 18S rRNA, guiding acetylation at C1773 (III in Figure 3D). Interestingly, structures of 90S particles show that Kre33, responsible for catalyzing both modifications (11,12), is bound to the 5’ domain of the 18S rRNA (Figure 3D, I), which is distant from helices h34 (II) and h45 (III) (37). However, Kre33 CRAC data revealed additional binding sites of Kre33 in the regions surrounding the acetylation sites (Figure 3D, Supplementary Figure S3 (11)), suggesting the occurrence of a yet structurally uncharacterized conformational rearrangement.

Mapping of the Sni445 CRAC reads on the snR4 and snR45 sequences revealed binding of Sni445 to both snoRNAs at their 3’ regions, near the conserved D boxes (Figure 3E-3H). Interestingly, these binding sites are 3’ of, but partially overlap with, those reported for Kre33 (Figure 3F, 3H).

### *SNR4* and *SNR45* deletions phenocopy the effects of *SNI445* deletion

Intriguingly, in the 40S subunit, Rps20, which physically and genetically interacts with Sni445 (Figure 1D and 1G), is in contact with helix h34 (Figure 4A, (38)). In particular, the positively charged residues R68 and K69, whose mutation resulted in a synergistic growth defect together with *SNI445* deletion (Figure 1G), are positioned in close proximity to ac4C1280 (Figure 4A). Notably, combining the *rps20.R68E/K69E* allele with deletion of *SNR4* also resulted in a synthetic growth defect (Figure 4B), suggesting that *RPS20* not only genetically interacts with *SNI445* but also with *SNR4*. In contrast, no genetic interaction was observed between *RPS20* and *SNR45* (Figure 4B).

**Figure 4.**
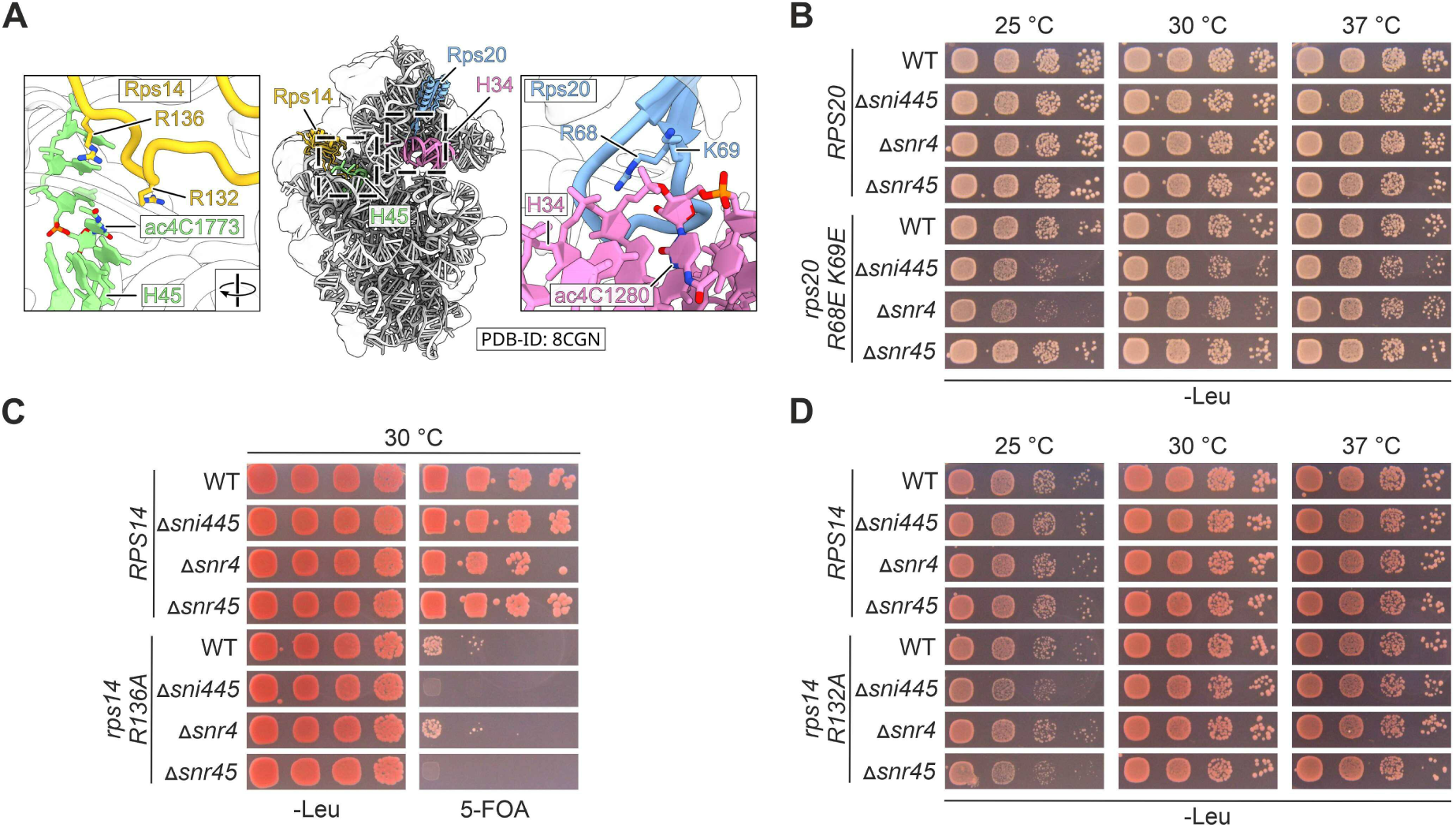
Deletion of *SNI445* is phenocopied by deletion of *SNR4* or *SNR45*. (A) Structure of the small ribosomal subunit (PDB 8CGN) (38). 18S rRNA (gray), Rps14 (light orange) and Rps20 (blue) are labelled and displayed as cartoon representation. 18S rRNA helices h34 and h45 are highlighted. The zoom-ins highlight the positions of ac4C1280, located near residues R68 and K69 of Rps20, and ac4C1773, located near residues R132 and R136 of Rps14. (B) *SNR4* genetically interacts with *RPS20*. The experiment was performed as described in Figure legend 1G, with the addition of *SNR4* or *SNR45* deletions. (C) Synthetic lethality between *rps14a.R136A* and deletion of *SNI445* or *SNR45*. Yeast cells (Δ*rps14a* Δ*rps14b*) carrying a *URA3* plasmid with wild-type *RPS14A* and *LEU2* plasmids with either wild-type *RPS14A* or the mutant *rps14a*.*R136A* allele, combined with *SNI445, SNR4,* or *SNR45* deletion, were spotted in serial dilutions on SDC-Leu and 5-FOA-containing plates and incubated for three days at 30°C. Lack of growth on 5-FOA plates indicates failure to lose the *URA3*-*RPS14A* plasmid, suggesting synthetic lethality. (D) Genetic enhancement of the mild growth defect of the *rps14.R132A* mutant upon deletion of *SNI445* or *SNR45*. Yeast cells (Δ*rps14a* Δ*rps14b*) carrying *LEU2* plasmids with either wild-type *RPS14* or mutant *rps14a*.*R132A* allele, combined with *SNI445, SNR4,* or *SNR45* knockout, were spotted in serial dilutions on SDC-Leu plates and incubated at 25°C, 30°C, or 37°C for two days.

In analogy to the genetic interaction between *SNR4* and *RPS20*, we asked whether a ribosomal protein is positioned near the C1773 acetylation site. Inspection of the 3D structure of the 40S subunit (38) revealed that the C-terminal tail of Rps14 is positioned in close proximity to ac4C1773 (Figure 4A). A previous study showed that mutation of the Rps14 C-terminal tail impairs cytoplasmic pre-40S maturation. In particular, when both *RPS14A* and *RPS14B* were deleted and complemented by a plasmid carrying *rps14b* alleles (39), replacement of R137 (corresponding to R136 in Rps14a) with alanine led to a strong accumulation of the 20S pre-rRNA (39). We tested a similar mutant, by introducing R136A into the more highly expressed *RPS14A* allele on a plasmid in a *rps14a*Δ *rps14b*Δ background. Strikingly, combining *sni445*Δ with this *rps14a.R136A* allele resulted in synthetic lethality (Figure 4C). Moreover, *SNR45* deletion caused synthetic lethality when combined with *rps14a.R136A*, mirroring the effect of *SNI445* deletion, whereas *SNR4* did not genetically interact with *rps14a.R136A* (Figure 4C). To validate these genetic interactions using a milder *rps14* mutant, we generated *rps14a.R132A,* which did not show a growth defect on its own (Figure 4D). Notably, both the *rps14.R132A sni445*Δ and the *rps14.R132A snr45*Δ double mutant displayed a synthetic growth defect at 25°C, while *SNR4* deletion did not affect growth of the *rps14.R132A* mutant (Figure 4D).

In summary, *SNR4* and *SNR45* deletion phenocopy the effects of *SNI445* deletion with respect to the genetic interactions with *RPS20* and *RPS14*. These results strongly suggest that the genetic interactions observed for *SNI445* arise from reduced recruitment and/or function of the acetylation guide snoRNAs snR4 and snR45.

### Sni445 interacts with acetyltransferase Kre33

Given that CRAC analyses did not reveal binding of Sni445 to rRNA, we hypothesized that Sni445 associates with 90S particles primarily through protein-protein interactions, potentially via Rps20 and Rps24, which showed a Y2H interaction with Sni445 (Figure 1C and 1D), possibly in conjunction with other proteins present in 90S particles.

To identify potential docking sites of Sni445, we performed TurboID-based proximity labeling (40,41). As expected, box C/D core proteins Nop56, Nop58 and Nop1 were enriched, with Nop56 showing the highest abundance and strongest enrichment. In contrast, Rps20 and Rps24, were not particularly enriched (Figure 5A). Among the prominently enriched ribosome assembly factors, there were factors associated with or acting on 90S/pre-40S particles, such as Kre33, Rrp12, Dhr2, and Rio1, but also pre-60S factors, including Erb1, Loc1, Nsa2, Rrp17, and Ssf1 (Figure 5A).

**Figure 5:**
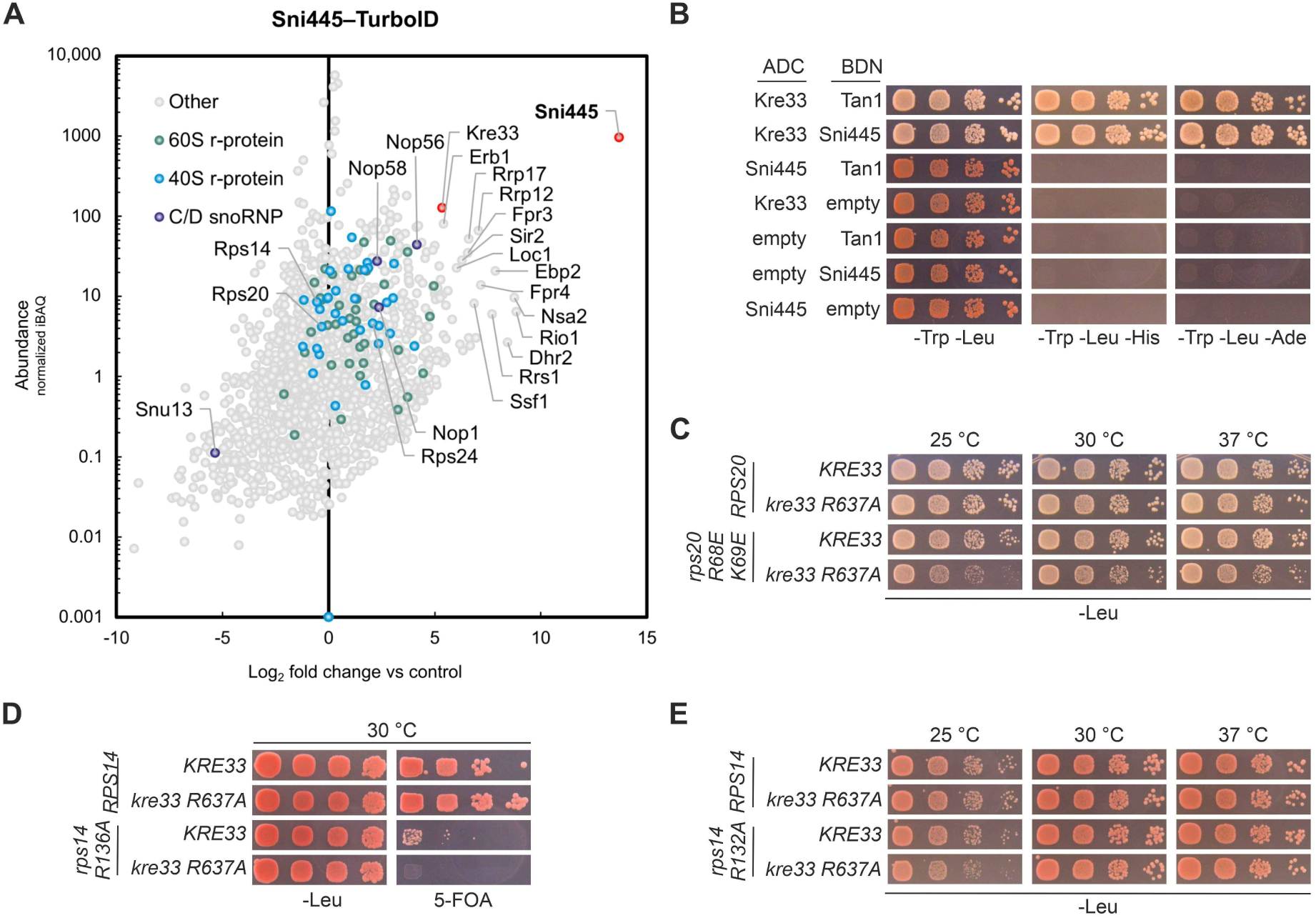
Sni445 interacts with the acetyltransferase Kre33. (A) Physical proximities of Sni445 revealed by TurboID-based proximity labelling. For each protein detected in the streptavidin pull-down of cells expressing Sni445-TurboID, the normalized abundance (iBAQ, intensity-based absolute quantification; y-axis) is plotted against its relative enrichment (log2-transformed; x-axis) compared to two negative control purifications (GFP-TurboID and NLS-GFP-TurboID baits, accounting for cytoplasmic and nuclear backgrounds, respectively). Proteins enriched relative to the controls can be found on the right side of the Christmas tree plot. (B) Y2H interaction assays between Sni445, Kre33 and the tRNA acetylation adaptor Tan1. Proteins were fused C-terminally to the Gal4 activation domain (ADC) or N-terminally to the Gal4 DNA binding domain (BDN). See Figure legend 1 (C-E) for details about media and growth conditions. (C) The catalytically inactive *kre33.R637A* mutant enhances the growth defect of the *rps20.R68E/K69E* mutant. Yeast cells (Δ*rps20*) carrying *LEU2* plasmids with either wild-type *RPS20* or the *rps20.R68E/K69E* allele in combination with wild-type *KRE33* or the genomically integrated *kre33.R637A* allele, encoding a catalytic inactive variant, were spotted in serial dilutions on SDC-Leu plates and incubated for three days at 25°C and for two days at 30°C and 37°C. (D) Catalytic inactivation of Kre33 (*kre33.R637A*) results in synthetic lethality with the *rps14a*.*R136A* mutation. Yeast cells (Δ*rps14a* Δ*rps14b*) carrying a *URA3* plasmid with wild-type *RPS14A,* and *LEU2* plasmids with either wildtype *RPS14A* or *rps14a.R136A* in combination with wild-type *KRE33* or the genomically integrated *kre33.R637A* allele were spotted in a serial dilution on SDC-Leu and 5-FOA containing plates and incubated for three days at 30°C. No growth on 5-FOA-containing plates indicates failure to lose the *URA3*-*RPS14A* plasmid, suggesting synthetic lethality. (E) The *kre33*.*R637A* mutation enhances the mild growth defect of the *rps14a*.*R132A* mutant. Yeast cells (Δ*rps14a* Δ*rps14b*) carrying *LEU2* plasmids with either wild-type *RPS14A* or the *rps14a.R132A* allele in combination with wild-type *KRE33* or the genomically integrated *kre33.R637A* allele were spotted in serial dilutions on SDC -Leu plates and incubated for two days at 25°C, 30°C, and 37°C.

The strong enrichment of Kre33, the acetyltransferase responsible for catalyzing the modifications guided by snR4 and snR45, together with the fact that both Sni445 (Figure 3A, B) and Kre33 (Figure 3F, H; (11)) interact with the snR4 and snR45 snoRNPs, suggested that Sni445 might directly interact with Kre33. Interestingly, Kre33 also modifies tRNAs, a process that is assisted by the adaptor protein Tan1 (15). Importantly, Y2H assays revealed that Sni445 strongly interacts with Kre33, to a similar extent as Tan1 (Figure 5B). In contrast, and as expected, Sni445 and Tan1 did not interact.

We next wondered if the phenotypes of *SNI445*, *SNR4* and *SNR45* deletion, apparent from their genetic interactions with *rps20* and *rps14* alleles, are the consequence of loss of acetylation at C1280 and C1773 or, alternatively, are due to a modification-independent function of snR4 and snR45. To distinguish between these two possibilities, we employed a catalytically inactive *kre33* mutant, in which the Kre33 protein carries an amino acid exchange at position 637 (R>A) (15). Notably, the *kre33.R637A* mutant displayed a synthetic growth defect when combined with the *rps20.R68E/K69E* allele (Figure 5C), thus recapitulating the effects of *SNI445* or *SNR4* deletion (Figure 4B). Similarly, synthetic lethality or a synthetic growth defect could be observed when the *kre33.R637A* allele was combined with the *rps14a.R136A* or, respectively, the *rps14a.R132A* allele (Figure 5D, 5E), phenocopying the impact of *SNI445* or *SNR45* deletion (Figure 4C, D). The observation that loss of the catalytic activity of Kre33 is sufficient to explain the synthetic growth defects is strong evidence that all observed genetic interactions result from loss of specific rRNA acetylation events.

### Sni445 recruits snR4 and snR45 to 90S particles to guide acetylation by Kre33

As Sni445 is a stable component of the snR4 and snR45 snoRNPs and interacts with the acetyltransferase Kre33, we next asked whether it is required for efficient acetylation at C1280 and C1773. To test this, we employed a previously established reverse transcription-based assay to detect N4-acetylcytidine (ac4C) (26). This method relies on the reduction of ac4C by sodium borohydride (NaBH_4_), resulting in conversion to N4-acetyl-3,4,5,6-tetrahydrocytidine. This modification causes reverse transcriptases to misincorporate nucleotides - most notably adenosine instead of guanosine - producing mixed C/T peaks in the sequencing readout following cDNA amplification. Consistent with previous findings (26,42), partial misincorporation was observed at both C1280 and C1773 in NaBH_4_-treated, but not in untreated wild-type samples (Figure 6A). In contrast, misincorporation was absent at C1280 in the *snr4*Δ and at C1773 in the *snr45*Δ strain, confirming loss of acetylation at these sites. Strikingly, both acetylation signals were fully abolished in the *sni445*Δ strain, indicating that Sni445 is essential for snR4 and snR45-guided cytidine acetylation (Figure 6A).

**Figure 6:**
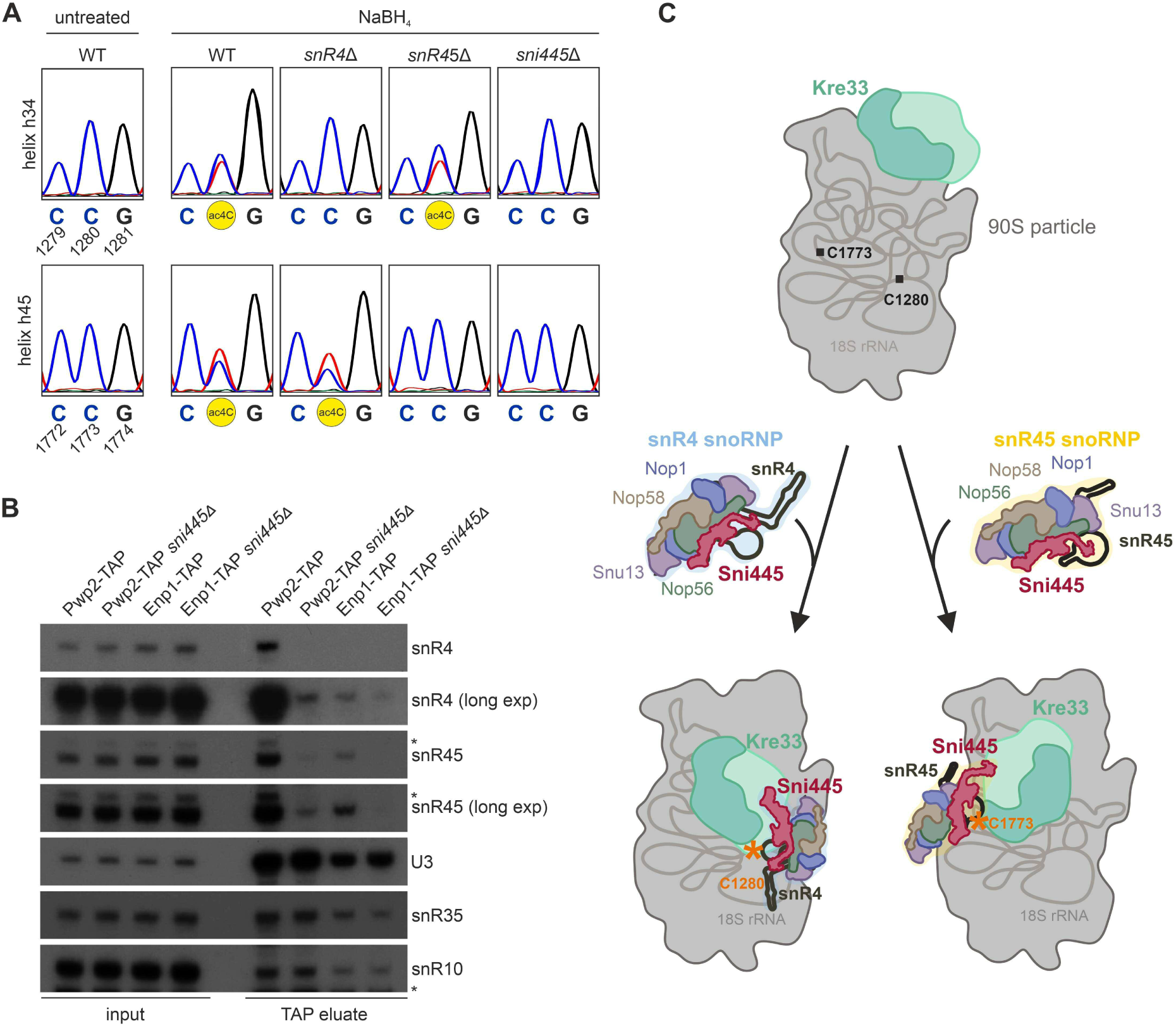
Sni445 recruits snR4 and snR45 to pre-ribosomal particles to facilitate acetylation by Kre33. (A) Sni445 is essential for acetylation at C1280 and C1773. Sanger sequencing analysis of PCR-amplified cDNAs generated from ac4C RNAs following NaBH4 treatment and reverse transcription. The red trace in the chromatograms represents Ts. Acetylation is apparent from a combined C/T peak in NaBH4-treated samples. (B) Deletion of *SNI445* alters the snoRNA composition of pre-ribosomal particles. RNA was extracted from TAP eluates of early 90S (Pwp2-TAP) and late 90S/pre-40S (Enp1-TAP) particles purified from wild-type and *sni445*Δ strains. Northern blot analysis was performed on inputs and TAP eluates using probes specific for snR4, snR45, U3, snR35, and snR10. Asterisks indicate residual signals from previously hybridized probes that were not fully removed during stripping. C. Model of the function of Sni445 in recruiting snR4 and snR45 to 90S particles to guide acetylation by Kre33.

Sni445 might either directly promote acetylation by Kre33 or act upstream by facilitating snR4 and snR45 snoRNP association with 90S particles. To distinguish between these possibilities, we affinity-purified early 90S particles using Pwp2-TAP and late 90S/pre-40S particles using Enp1-TAP as baits and assessed the effects of *SNI445* deletion on the association of snR4 and snR45 by northern blotting (Figure 6B). Importantly, the overall expression and stability of snR4 and snR45 were unaffected by *SNI445* deletion, as shown by the input controls. In wild-type cells, both snoRNAs were detected in Pwp2-purified particles, and to a lesser extent, in Enp1-purified particles (Figure 6B). However, in the *sni445*Δ strains, the association of snR4 and snR45 with these pre-ribosomal particles was substantially reduced, whereas the binding of other snoRNAs (i.e. U3, snR35, and snR10) remained unchanged (Figure 6B). These findings suggest that Sni445 promotes the recruitment or stabilization of snR4 and snR45 within 90S particles, thereby enabling Kre33 to catalyze acetylation at C1280 and C1773 (Figure 6C).

## Discussion

In this study, we identify Sni445 as a novel ribosome assembly factor and a previously unrecognized auxiliary component of the snR4 and snR45 box C/D snoRNPs. Unlike canonical box C/D snoRNPs, which guide 2’-O methylation, snR4 and snR45 direct acetylation of 18S rRNA residues C1280 and C1773, respectively, via the acetyltransferase Kre33 (11,12). Our data show that Sni445 is associated with free snR4 and snR45 snoRNPs (Figure 3C). Interestingly, although Kre33 crosslinks to both snoRNAs (11), and directly interacts with Sni445 (Figure 5B), it is not a stoichiometric component of Sni445- and Nop58-containing snoRNP complexes (Figure 3C), suggesting that Kre33 interacts transiently with these snoRNPs or joins them within 90S particles through a direct interaction with Sni445.

Notably, Kre33 is an essential structural component of the 90S particle (43–45), whereas snR4, snR45, and Sni445 are non-essential, suggesting that Kre33 must associate with 90S particles independent of the snR4 and snR45 snoRNPs. Intriguingly, in cryo-EM structures of 90S particles, Kre33 is positioned far from its acetylation target sites, C1280 and C1773 (Figure 3D, (37,45–47)). The 90S structural state in which snR4 and snR45 are bound has not yet been visualized by cryo-EM, but some form of rRNA rearrangement or Kre33 positioning must occur to bring these target nucleotides into vicinity of the Kre33 catalytic center. It is likely that the snR4 and snR45 snoRNPs, with Sni445 as key player, might be involved in this task (Figure 6C).

Previous CRAC analyses revealed that Kre33 binds three distinct regions of the 18S rRNA (11), all of which are somehow connected to Sni445: Kre33 binds helices h33-h36 in the 3’ major domain, the site where snR4 guides acetylation and where the internal loop of Rps20 is located (Figure 3D, 4A, Supplementary Figure S3). Additionally, Kre33 interacts with helices h44-h45 in the 3’ minor domain, which include the snR45 target region and lie adjacent to the C-terminal extension of Rps14 (11) (Figure 3D, 4A, Supplementary Figure S3). The third Kre33-binding site encompasses helices h7-h10 in the 5’ domain, near the Rps24-binding site (11) (Figure 3D, Supplementary Figure S3). The observed Y2H interaction between Sni445 and Rps24 (Figure 1C), alongside Kre33 binding near this region, suggests an additional, yet unexplored function shared by Sni445 and Kre33.

Strikingly, deletion of Sni445 abolishes acetylation at C1280 and C1773 (Figure 6A) and dramatically reduces the association of snR4 and snR45 with 90S particles (Figure 6B). This strongly suggests that Sni445 is required for the recruitment and/or stable anchoring of these snoRNPs to 90S particles. This finding challenges the prevailing view that snoRNA guide sequences are sufficient to mediate targeting, and implies that, at least for snR4 and snR45, additional recruitment mechanisms involving Sni445 are required.

Our previous work has shown that the *rps20*.*R68E/K69E* internal loop mutant is impaired in cytoplasmic 40S maturation due to reduced Rio2 ATPase activation and release (28), while the Woolford laboratory demonstrated that mutations within the C-terminal extension of Rps14 (e.g., R136A) compromise cytoplasmic cleavage of the 20S pre-rRNA to mature 18S rRNA (39). The observed genetic interactions between *SNI445*/*SNR4*/*SNR45* and these ribosome biogenesis mutants suggest that rRNA acetylation mediated by Kre33 may contribute not only to ribosome function but also to ribosome assembly.

Beyond the Sni445-containing complexes studied in our work, our data suggest that Sni445 establishes additional interactions: Our TAP- and FLAG-purification experiments identified pre-60S assembly factors co-enriched with Sni445 (Figure 2C), and the TurboID-based proximity labeling experiment further supports a physical proximity to pre-60S factors (Figure 5A). Moreover, snR4 was previously shown to bind 5S rRNA (nt 22–32) and 25S rRNA (nt 1866–1878) (48). These data raise the possibility that Sni445, perhaps as part of snR4/snR45 snoRNPs, might also participate in pre-60S biogenesis. Interestingly, FLAG-purification data suggest that Sni445 might associate with the Paf complex and RNase H2. The known role of the Paf complex in snoRNA 3’ end formation (31,32) raises the possibility that Sni445 is recruited to these snoRNAs already co-transcriptionally. The functional characterization of these potential interactions of Sni445 with the Paf complex and with pre-60S particles would be interesting subjects for future studies.

Interestingly, while canonical snoRNPs only contain four different core proteins, there are a few examples of atypical snoRNPs containing distinct additional protein components. For instance, Rrp9 is a component of the box C/D U3 snoRNP, Utp23, Kri1 and Krr1 of the box H/ACA snR30 snoRNP, while the Npa1 complex is associated with the box C/D snR190 snoRNP (13,14,49–51). We now add Sni445 to this list of dedicated components of snoRNPs. Of note, all these atypical snoRNPs are involved in functions other than the canonical roles of their snoRNA classes in guiding methylation or pseudouridylation.

## Data availability statements

The MS proteomics raw data together with the processing files were deposited to the ProteomeXchange Consortium using the PRIDE partner repository (https://www.ebi.ac.uk/pride/) with the dataset identifier PXD065447. Reviewers can access the dataset by log in to the PRIDE website using the following details: Project accession: PXD065447; Token: vzPAtkd6lx9E. NGS analysis files of raw and processed data were deposited in the Gene Expression Omnibus database under the accession number GSE299803 (For review only: secure token: ebkbqmkyjjclfon). TurboID raw and processed data are provided in Supplementary Table S3.

## Acknowledgment

We are grateful to Virginie Marchand and Yuri Motorin (EpiRNA-Seq, IBSLor, Nancy, France) for the sequencing of CRAC libraries. We thank members of the Henry/Henras, Pertschy, Bergler and Mitterer groups for helpful discussion.

## Author contribution statement

BP, JH, IZ, and AKH conceived the study. JH, IZ, HH, SF, MT, NK, SR, and BP performed the experiments, and JH, IZ, HH, SF, MT, NK, RB, US, DK, AKH, and BP analyzed the data. HH, SF, BA, TC, MA, and US contributed to bioinformatics analysis. JH, BP, DK, and AKH wrote the manuscript with input from all authors.

## Footnotes

1 In this manuscript, r-proteins are referred to by their standard names as listed in the *Saccharomyces* Genome Database, SGD, www.yeastgenome.org). Upon first mention of each r-protein, the corresponding name from the recently introduced new nomenclature (18) is provided in parentheses.

