## Supplementary Data for "Sni445 recruits box C/D snoRNPs snR4 and snR45 to guide ribosomal RNA acetylation by Kre33"

### Supplementary Figures

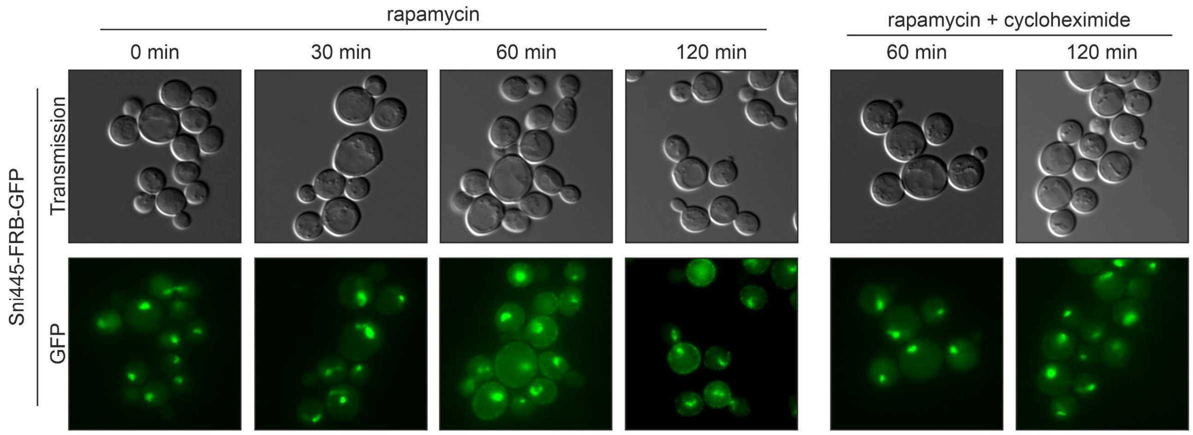


**Supplementary Figure S1: Sni445 localizes exclusively to the nucleus**. Anchor-away yeast reporter cells (1) expressing Sni445-FRB-GFP were treated during logarithmic growth phase with 1 µg/ml rapamycin or with 1 µg/ml rapamycin plus 10 µg/ml cycloheximide (to inhibit de-novo protein synthesis), followed by fluorescence microscopy. Samples were collected after 30, 60, and 120 min of rapamycin treatment, or after 60 and 120 min of combined rapamycin and cycloheximide treatment.

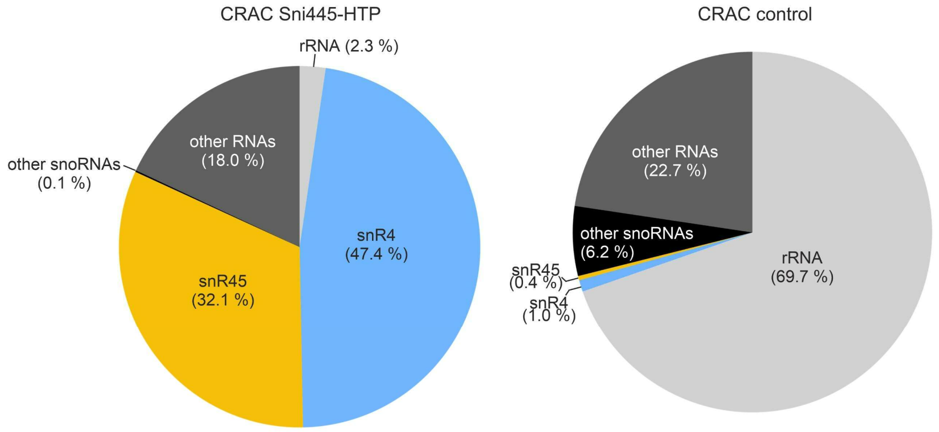


**Supplementary Figure S2: Sni445 CRAC analysis**. Pie charts showing the distribution of RNA classes identified in CRAC experiments, comparing Sni445-HTP (left panel) to an untagged negative control strain (right panel).

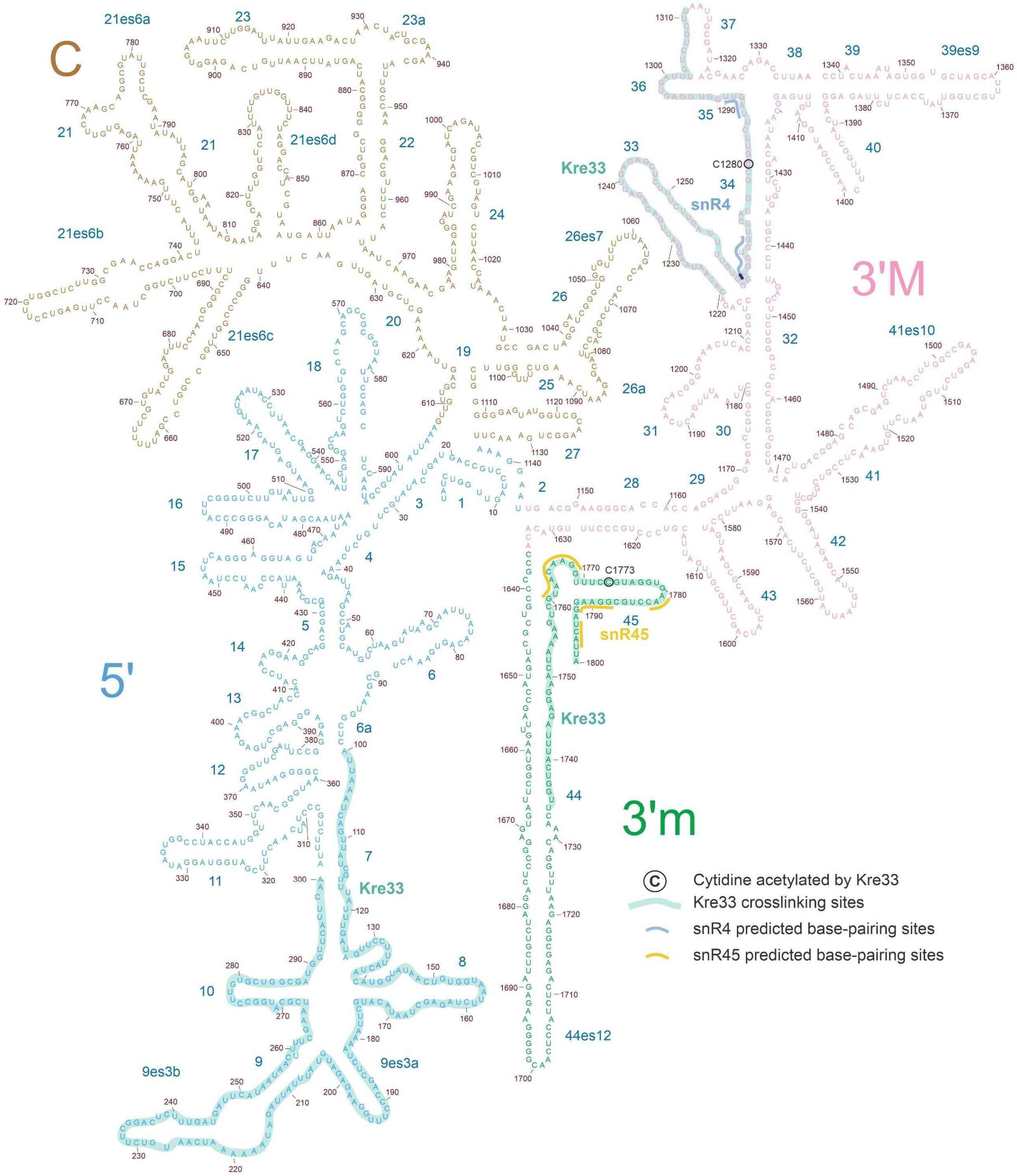


**Supplementary Figure S3:** Secondary structure of 18S rRNA. The four secondary structure domains, 5’ domain (5’; blue), central domain (C; brown), 3’ major domain (3’M; pink), and 3’ minor domain (3’m; green) are highlighted. Base-pairing sites of snR4 (blue) and snR45 (yellow), as well as crosslinking sites of Kre33 (turquoise) are highlighted (2). Cytidines acetylated by Kre33 are indicated by circles.

**Supplementary Tables**

**Supplementary Table S1. Yeast strains used in this study**

| name | genotype | source |
|---|---|---|
| W303 | *ade2 leu2 his3 trp1 ura3* | (3) |
| W303 *ade3*Δ | *MAT*α *ade2 leu2 his3 trp1 ura3 ade3*Δ::kanMX4 | (4) |
| W303 *ade3*Δ | *MAT*α *ade2 leu2 his3 trp1 ura3 ade3*Δ::natNT2 | This study |
| Y2H PJ69-4A | *MAT***a** *trp1-901 leu2-3,112 ura3-52 his3-200 gal4*Δ *gal80*Δ *LYS2::GAL1-HIS3 GAL2-ADE2 met2::GAL7*-lacZ | (5) |
| Sni445-GFP Nop58-RedStar2 | W303 *MAT***a** *NOP58*-RedStar2*::natNT2, SNI445*-GFP*::*HIS3MX6 | This study |
| Sni445 -FRB-GFP | *MAT*α *leu2 ura3 his3 ade2 tor1-1 fpr1::*natNT2 *PMA1*-2xFKBP12*::TRP1 SNI445*-FRB-GFP*::*kanMX | This study |
| BY4741 | *MAT***a** *his3 leu2 met15 ura3* | Euroscarf |
| Sni445-HTP | BY4741 *MAT***a** *SNI445-HTP::*klURA3 | This study |
| Sni445-TAP | W303 *MAT***a** *SNI445*-TAP*::*HIS3MX6 | This study |
| Sni445-FLAG | W303 *MAT*α *SNI445*-FLAG*::*natNT2 | This study |
| Pwp2-TAP | W303 *MAT***a** *PWP2*-TAP::HIS3MX6 | This study |
| Pwp2-TAP *sni445*Δ | W303 *MAT***a** *PWP2*-TAP*::*HIS3MX6 *sni445*Δ*::*kanMX | This study |
| Enp1-TAP | W303 *MAT***a** *ENP1*-TAP*::*HIS3MX6 | (6) |
| Enp1-TAP s*ni445*Δ | W303 *MAT***a** *ENP1*-TAP*::*HIS3MX6 *sni445*Δ*::*kanMX4 | This study |
| *RPS20*-Shuffle | W303 *MAT*α *rps20*Δ*::*HIS3MX4 *ade3*Δ*::*kanMX4 [YCplac33-*RPS20*] | This study |
| *RPS20*-Shuffle *sni445*Δ | W303 *MAT*α *rps20*Δ*::*HIS3MX4 *ade3*Δ*::*kanMX4 [YCplac33-*RPS20*] *sni445*Δ*::*natNT2 | This study |
| *RPS20*-Shuffle *snr4*Δ | W303 *MAT*α *rps20*Δ*::*HIS3MX4 *ade3*Δ*::*kanMX4 [YCplac33-*RPS20*] *snr4*Δ*::*natNT2 | This study |
| *RPS20*-Shuffle *snr45*Δ | W303 *MAT*α *rps20*Δ*::*HIS3MX4 *ade3*Δ*::*kanMX4 [YCplac33-*RPS20*] *snr45*Δ*::*natNT2 | This study |
| *RPS14*-Shuffle | W303 *MAT***a** *rps14a*Δ*::*HIS3MX6 *rps14b*Δ*::*natNT2 [YCplac33-*RPS14A*] | This study |

| | | |
|---|---|---|
| *RPS14*-Shuffle *sni445*Δ | W303 *MAT***a** *rps14a*Δ*::*HIS3MX6 *rps14b*Δ*::*natNT2 *sni445*Δ*::*kanMX4 [YCplac33-*RPS14A*] | This study |
| *RPS14*-Shuffle *snr4*Δ | W303 *MAT***a** *rps14a*Δ*::*HIS3MX6 *rps14b*Δ*::*natNT2 *snr4*Δ*::*kanMX4 [YCplac33-*RPS14A*] | This study |
| *RPS14*-Shuffle *snr45*Δ | W303 *MAT***a** *rps14a*Δ*::*HIS3MX6 *rps14b*Δ*::*natNT2 *sni45*Δ*::*kanMX4 [YCplac33-*RPS14A*] | This study |
| *RPS14*-Shuffle *KRE33* WT | W303 *MAT***a** *rps14a*Δ*::*HIS3MX6 *rps14b*Δ*::*natNT2 [YCplac33-*RPS14A*] *KRE33::*hphNT1 | This study |
| *RPS14*-Shuffle *kre33.R637A* | W303 *MAT***a** *rps14a*Δ*::*HIS3MX6 *rps14b*Δ*::*natNT2 [YCplac33-*RPS14A*] *kre33.R637A::*hphNT1 | This study |
| *RPS20*-Shuffle *KRE33 WT* | W303 *MAT*α *rps20*Δ*::*HIS3MX4 *ade3*Δ*::*kanMX4 [YCplac33-*RPS20] KRE33::*hphNT1 | This study |
| *RPS20*-Shuffle *kre33.R637A* | W303 *MAT*α *rps20*Δ*::*HIS3MX4 *ade3::*ΔkanMX4 [YCplac33-*RPS20*] *kre33.R637A::*hphNT1 | This study |
| *sni445*Δ | W303 *MAT*α *ade3*Δ::natNT2 *sni445*Δ::klTRP1 | This study |
| *rps24b*Δ | W303 *MAT*α *ade3*Δ::natNT2 *rps24b*Δ::kanMX | This study |
| *rps24b*Δ *sni445*Δ | W303 *MAT*α *ade3*Δ::natNT2 Δ*rps24b*Δ::kanMX6 *sni445*Δ::klTRP1 | This study |
| *rps24a*Δ | W303 *MAT*α *ade3*Δ::natNT2 Δ*rps24a*Δ::kanMX6 | This study |
| *rps24a*Δ *sni445*Δ | W303 *MAT*α *ade3*Δ::natNT2 Δ*rps24a*Δ::kanMX6 *sni445*Δ::klTRP1 | This study |
| *RPS2* shuffle | W303 *MAT***a** *rps2*Δ::kanMX4 [pRS316-*RPS2*] | (7) |
| *RPS2* shuffle *sni445*Δ | W303 *MAT***a** *rps2*Δ::kanMX4 [pRS316-*RPS2*] *sni445*ΔHIS3MX6 | This study |
| *sni445*Δ | W303 *MAT*α *sni445*Δ:: klTRP1 | This study |
| *snr4*Δ | W303 *MAT*α *snr4*Δ::kanMX6 | This study |
| *snr45*Δ | W303 *MAT*α *snr45*Δ:: hphNT1 | This study |
| Sni445-TAP Nop58-FLAG | W303 *MAT*α *SNI445*-TAP::klURA3 *NOP58*-FLAG::natNT2 | This study |

**Supplementary Table S2. Plasmids used in this study**

| **name** | **genotype** | **source** |
|---|---|---|
| pFA6a-HIS3MX4 | for chromosomal deletion | (8) |
| pFA6a-kanMX4 | for chromosomal deletion | (8) |

| | | |
|---|---|---|
| pFA6a-natNT2 | for chromosomal deletion | (9) |
| pFA6a-klTRP1 | for chromosomal deletion | (10) |
| pFA6a-hphNT1 | for chromosomal deletion | (9) |
| pFA6a-*KRE33*(1800-Stop)-150 downstream of *KRE33* - hphNT1 - 226-682 downstream *KRE33* | for chromosomal insertion of *KRE33* wild-type sequence | This study |
| pFA6a- *kre33 R636A*(1800-Stop)-150 downstream of *KRE33* - hphNT1 - 226-682 downstream *KRE33* | for chromosomal insertion of *kre33 R636A* mutant sequence | This study |
| pFA6a-GFP(S65T)::HIS3MX4 | for C-terminal tagging | (11) |
| pFA6a-TAP::HIS3MX4 | for C-terminal tagging | (12) |
| pBS1539 HTP::klURA3 | for C-terminal tagging | (13) |
| pFA6a-FRB-GFP::kanMX | for C-terminal FRB-GFP tagging | (1) |
| pFA6a Flag-TCYC1-natNT2 | for C-terminal tagging | (14) |
| pCUP111-yEGFP-(GA)5-TurboID-2xHA (pDK9295) | *CEN, LEU2,* P*CUP1, yEGFP*-TurboID-2xHA | (15) |
| pCUP111-SV40NLS-yEGFP-(GA)5-TurboID-2xHA (pDK9296) | *CEN, LEU2,* P*CUP1, SV40NLS*-yEGFP-TurboID-2xHA, T*ADH1* | (16) |
| pCUP111-*SNI445*-(GA)5-TurboID-2xHA | *CEN, LEU2,* P*CUP1, SNI445*-TurboID-2xHA, T*ADH1* | This study |
| pGAG4ADC111-*RPS14A* | CEN, *LEU2,* P*ADH1,* T*ADH1*, C-terminal $(GA)_5$-G4AD-HA | (7) |
| pGAG4ADC111-*SNI445* | CEN, *LEU2,* P*ADH1,* T*ADH1*, C-terminal $(GA)_5$-G4AD-HA | This study |
| pGAG4ADC111-*RPS10* | CEN, *LEU2,* P*ADH1,* T*ADH1*, C-terminal $(GA)_5$-G4AD-HA | This study |
| pG4BDN22-*SNI445* | CEN, *TRP1,* P*ADH1,* T*ADH1*, N-terminal G4BD-cMyc | This study |
| pG4BDN22-*RPS20* | CEN, *TRP1,* P*ADH1,* T*ADH1*, N-terminal G4BD-cMyc | This study |
| pGAG4ADC111-*KRE33* | CEN, *LEU2,* P*ADH1,* T*ADH1*, C-terminal $(GA)_5$-G4AD-HA | This study |

| | | |
|---|---|---|
| pG4BDN22-*TAN1* | CEN, *TRP1,* P*ADH1,* T*ADH1*, N-terminal G4BD-cMyc | This study |
| YCplac111-*RPS20* | *CEN*, *LEU2*, *RPS20* | (17) |
| YCplac111-*rps20.R68E/K69E* | *CEN*, *LEU2*, *rps20 R68/K69>E* | (17) |
| YCplac111-*RPS14A* | *CEN*, *LEU2*, *RPS14A* | (18) |
| YCplac111-*rps14a.R132A* | *CEN*, *LEU2*, *rps14a.R132A* | This study |
| YCplac111-*rps14a.R136A* | *CEN*, *LEU2*, *rps14a.R136A* | (18) |
| pRS314 *RPS2* | *CEN*, *TRP1*, *RPS2* | (7) |
| pRS314 *rps2-1* | *CEN*, *TRP1*, *rps2-1* | (7) |

Unless otherwise stated, all genes were cloned with their natural promoters.

**Supplementary Table S3. Sni445 TurboID raw and processed data.**

The table is provided separately as an Excel file.
